# Control-based continuation: a new approach to prototype synthetic gene networks

**DOI:** 10.1101/2021.12.21.473142

**Authors:** Irene de Cesare, Davide Salzano, Mario di Bernardo, Ludovic Renson, Lucia Marucci

## Abstract

Control-Based Continuation (CBC) is a general and systematic method to carry out the bifurcation analysis of physical experiments. CBC does not rely on a mathematical model and thus overcomes the uncertainty introduced when identifying bifurcation curves indirectly through modelling and parameter estimation. We demonstrate, *in silico*, CBC applicability to biochemical processes by tracking the equilibrium curve of a toggle switch which includes additive process noise and exhibits bistability. We compare results obtained when CBC uses a model-free and model-based control strategy and show that both can track stable and unstable solutions, revealing bistability. We then demonstrate CBC in conditions more representative of a real experiment using an agent-based simulator describing cells growth and division, cell-to-cell variability, spatial distribution, and diffusion of chemicals. We further show how the identified curves can be used for parameter estimation and discuss how CBC can significantly accelerate the prototyping of synthetic gene regulatory networks.

## Introduction

The complexity of biological systems is widely acknowledged. In native organisms, multi-scale intracellular interactions often result in complex nonlinear dynamics. Consider, for example, switches and oscillations in gene expression, which are used by cells to process external inputs and program specific cell outputs. Synthetic biology aims at engineering biological computation by recreating such dynamics using circuits embedded into living cells [1, 2, 3, 4, 5, 6, 7, 8, 9, 10].

Mathematical modelling is widely used within synthetic biology design-build-test-learn cycles. In the context of engineered gene regulatory networks, models can both support the design phase (indicating the parameter space which allows the emergence of the desired dynamics, such as oscillations), and the testing upon experimental implementation. Moreover, modelling is currently the only way in synthetic biology to study the relationship between physical parameters variations and bifurcations; the latter represent stability boundaries where qualitative and quantitative changes to the system’s dynamics occur. For instance, saddle-node and Hopf bifurcations (responsible for hysteresis and oscillatory behaviours, respectively) that are commonly observed in native biological systems [11] can only be studied by nonlinear model analysis. This means that, firstly, a mathematical model needs to be derived and fitted to often *ad hoc* generated experimental data. Then, nonlinear model analysis can be performed to identify the bifurcation behaviour being observed.

The derivation of biochemical models can however be challenging, both in terms of model structure (which depends on underlying hypothesis on the system), and parameter identification and validation (which can be troublesome in the case of incomplete/noisy experimental datasets). Model uncertainties will inevitably result in misleading conclusions [12, 13]. As a consequence, the design and testing of engineering synthetic biochemical circuits that perform as intended is extremely difficult, unless various design, build, test and learn iterations are performed [14].

Control-based continuation (CBC), originally proposed by Sieber and Krauskopf [15], is a general, systematic and model-free testing method that applies the principles of numerical continuation (a computational method for bifurcation analysis) to physical experiments. The fundamental principles of CBC are well established and the method has been applied to a wide range of mechanical systems. For instance, Barton et al. [16] studied the periodic responses of a bistable energy harvester. A similar system was studied by Renson et al. [17]. The continuation method was also demonstrated on a multi-degree-of-freedom structure exhibiting isolated curves of periodic responses and quasi-periodic oscillations [18]. CBC exploits feedback control to explore the nonlinear dynamics of a system, detect bifurcations, and eventually trace their evolution with respect to adjustable parameters directly during experimental tests.

Recently, external feedback controllers have been exploited for controlling gene expression in living cells by means of microfluidics/microscopy or optogenetics platforms [19, 20, 21, 22, 23, 24, 25, 26, 27]. From a control theory standpoint, genetic networks are the processes to be controlled, while the controller is implemented on a computer. Biosensors are used to measure the state of the process, usually by means of fluorescent reporter proteins. The fluorescence evaluation can be done either at single-cell or at cell population level, and is fed to a control algorithm that evaluates the appropriate control input aiming to steer the process to a selected reference signal. Inputs are then actuated on the process via light or chemical compounds [28].

By employing external feedback controllers to steer gene expression it should be possible to apply CBC to track nonlinear dynamics in biochemical systems, overcoming the need for a model identification step and defining a shorter way for bifurcation tracking which includes the isolation of unstable equilibria (see in Fig. 1A path 1 compared to path 2).

**Figure 1:**
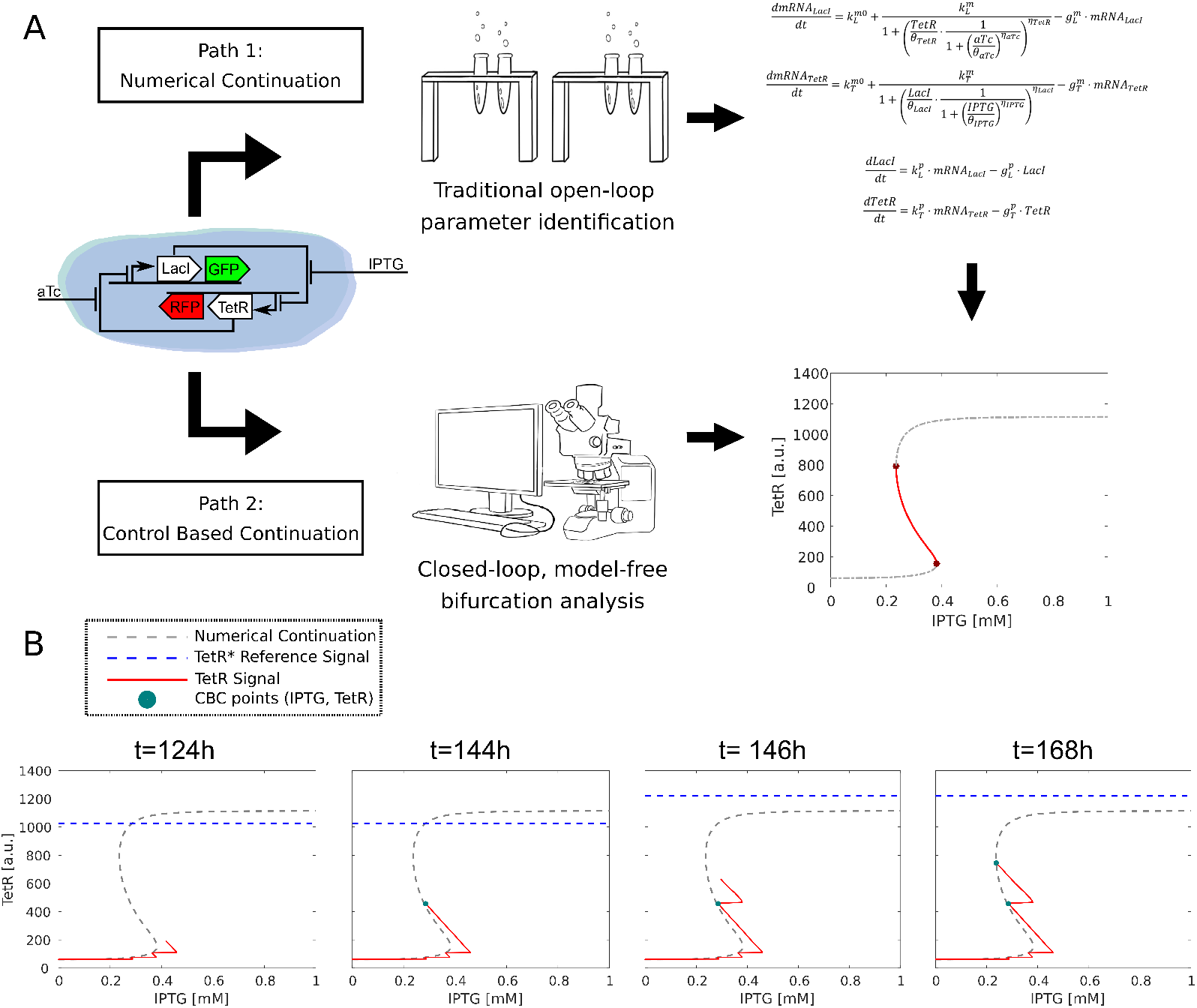
A) A schematic Toggle Switch from Lugagne et al. [6]; two different paths to the bifurcation diagram (- -): numerical continuation (path 1), and control-based continuation (path 2). Unstable branch of the bifurcation curve (−) and bifurcation points (*) are highlighted. B) Steps of the CBC algorithm: at time *t* = 124*h* the control pushes the system’s trajectory *TetR*(*IPTG*) (–) towards the new reference *TetR\** (- -). At time *t* = 144*h* transient dynamics are extinguished and a point 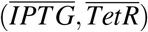 is saved (.). Then, the reference signal is increased and the process is repeated.

Here, we demonstrate, for the first time, the applicability of CBC to prototype the dynamics of a synthetic gene regulatory network and to estimate the parameters of a fixed model structure, if available, using data collected after the CBC routine. We use as a test bed the toggle switch, a bistable biological circuit, firstly embedded in *Escherichia coli* cells by Gardner et al. [10], which is often used to benchmark new control strategies and is considered a fundamental tool for cell computation [11, 29]. Specifically, we run *in silico* experiments on a recent version of the toggle switch, for which external feedback control was shown to be successful [6]. Moreover, we leveraged an agent-based simulator called BSim [30] to test the performance of the designed algorithms in conditions more representative of a real experiment. Our *in silico* demonstration of CBC gives us the confidence that the method should be exploitable *in vivo* to fully explore the parameter space which allows the emergence of the desired dynamics (in our example, hysteresis), leading to rapid and model-free characterization, and also parameter estimation, of engineered synthetic modules.

## Results and Discussion

CBC retrieves stable and unstable invariant solutions of a dynamical system through the application of an external control action. In order to acquire the bifurcation diagram, the controller must not modify the position in the parameter space of the uncontrolled system’s invariant solutions. For example, the equilibria of a controlled system are in general different from the one of the uncontrolled system and, to recover the response of the underlying uncontrolled system of interest, CBC seeks a control signal that decays asymptotically to zero, i.e.,

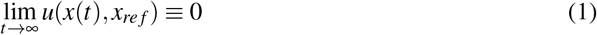

where *x* ∈ ℝ is the system’s state, *x_ref_* ∈ ℝ is the control reference signal and *t* is time. Although the control signal is asymptotically converging to zero, it is synthetized in order to stabilise local dynamics of the system such that unstable equilibria become stable and hence detectable forward in time. A control strategy that satisfies (1) is called noninvasive and does not modify the system’s equilibrium positions in the parameter space. Finding a noninvasive control signal usually requires finding the “right” reference input (*x_ref_*) for the controller such that (1) is satisfied, while the solution of interest is stabilised.

When the control input can be chosen as the bifurcation parameter of interest, the methodology can be significantly simplified as the control signal is only required to settle to a constant value. Indeed, when this condition is achieved, the non-zero constant control signal can be viewed as a mere shift in the bifurcation parameter value. This simplified method is used in this paper, and more details about its implementation can be found in the Methods section. Furthermore, CBC can also be extended to characterise systems exhibiting oscillations and other bifurcations as described in [17, 18, 31, 32].

The applicability of CBC is demonstrated here in *silico* on a synthetic gene network, the toggle switch (Fig. 1). The mathematical model developed by Lugagne et al. [6] is used. From a control system standpoint, it can be seen as a 2-input (*aTc* and *IPTG*) 2-output (*LacI* and *TetR*) nonlinear dynamical system. The inputs correspond to drugs that can be added to the cells environment, while the outputs are proteins synthesized by cells and detected by tagging them with fluorescent reporters. Here the control signal (specifically *IPTG*) plays a dual role: not only it pushes the system into a new state, but it keeps the system stable allowing it to explore the nearby unstable dynamics which would not be collected otherwise (see the red branch of the bifurcation diagram in Fig. 1 A). The model derivation and further information about the network can be found in the Methods section.

Fig. 1 B illustrates an experiment conducted with CBC; data points collected with CBC (blue point) are compared to the actual bifurcation curve obtained using standard model-based numerical continuation algorithms (gray dashed line). First, a particular control reference value (*x_ref_* = *TetR*\*) is selected (dashed-blue line). The reference signal is compared to the actual expression of the gene of interest to compute a control error. This is fed to the controller, which evaluates a controlling signal (*IPTG* concentration), translated into an input to the toggle switch that will therefore change its state (the red line in Fig. 1 B shows the system transient dynamic). At every sampling time a new measure is acquired and the entire process is repeated until a steady state value is reached. The steady state value of the state variables, together with the associated control signal, are then saved to define a point 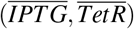 in the bifurcation diagram. Subsequently, a new reference value is picked (see Methods) and the entire process is repeated. To trace out the entire equilibrium curve, the continuation algorithm requires a set of control reference values, broadly covering the range of gene expression of interest (in our case the *TetR* fluorescence expression).

CBC is a model-free method because it does not require the knowledge of a mathematical model of the system to be applied to, and the results’ accuracy does not depend on the structure and parameter values of a model. Furthermore, the controller within CBC is only required to be stabilizing and noninvasive. As such, CBC is not tied to a specific type of feedback controller and control laws that do not require a mathematical model of the system can be used. However, the design, parameter tuning and overall performance of the controller can be improved if some knowledge (like a model) of the system dynamics is available. Here, the use of a model-free proportional (P) controller and a model-predictive controller (MPC) is compared. The former controller was chosen as it is widely used in CBC applications, while the latter is commonly exploited in synthetic biology [20]. When using Model Predictive control strategies, simple linear models work well, showing that detailed mathematical representations are not needed for CBC to be work. A sampling time of 5 minutes is considered in this work; this is a realistic time interval that we previously used to image and externally control bacterial cells [22, 25]. Note that, with the simplified CBC approach, the controller is not required to take the control error to zero but only to stabilise the system to the nearest equilibrium point. This does not affect the accuracy of results, as further explained in Methods.

### CBC can reconstruct the toggle switch bifurcation diagram under deterministic simulation settings

We first applied CBC on the deterministic model of the toggle switch. During the in *silico* experiment, 30 steady state points 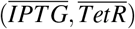 were acquired by varying the reference input to the control strategy between *TetR*\* = 1800[*a.u.*] (or 1200[*a.u.*]) and *TetR*\* = 0[*a.u*]. The maximum duration of a single equilibrium acquisition was constrained to 9 hours and 55 minutes, after which the control reference was modified. The steady states values were computed by taking the average of the samples recorded in the last 60 minutes (12 samples) of the simulation carried out for each value of the reference input. Averaging is not essential in a deterministic scenario, but becomes fundamental in a noisy environment as a way to filter out some noise. Points collected with the CBC perfectly overlap with the numerical continuation diagram (Fig. 2 A and 3 A). Both control strategies have comparable tracking performances and proved able to stabilise the unstable branch of equilibria delimited by the Saddle-Node bifurcations, capturing the whole bifurcation curve.

**Figure 2:**
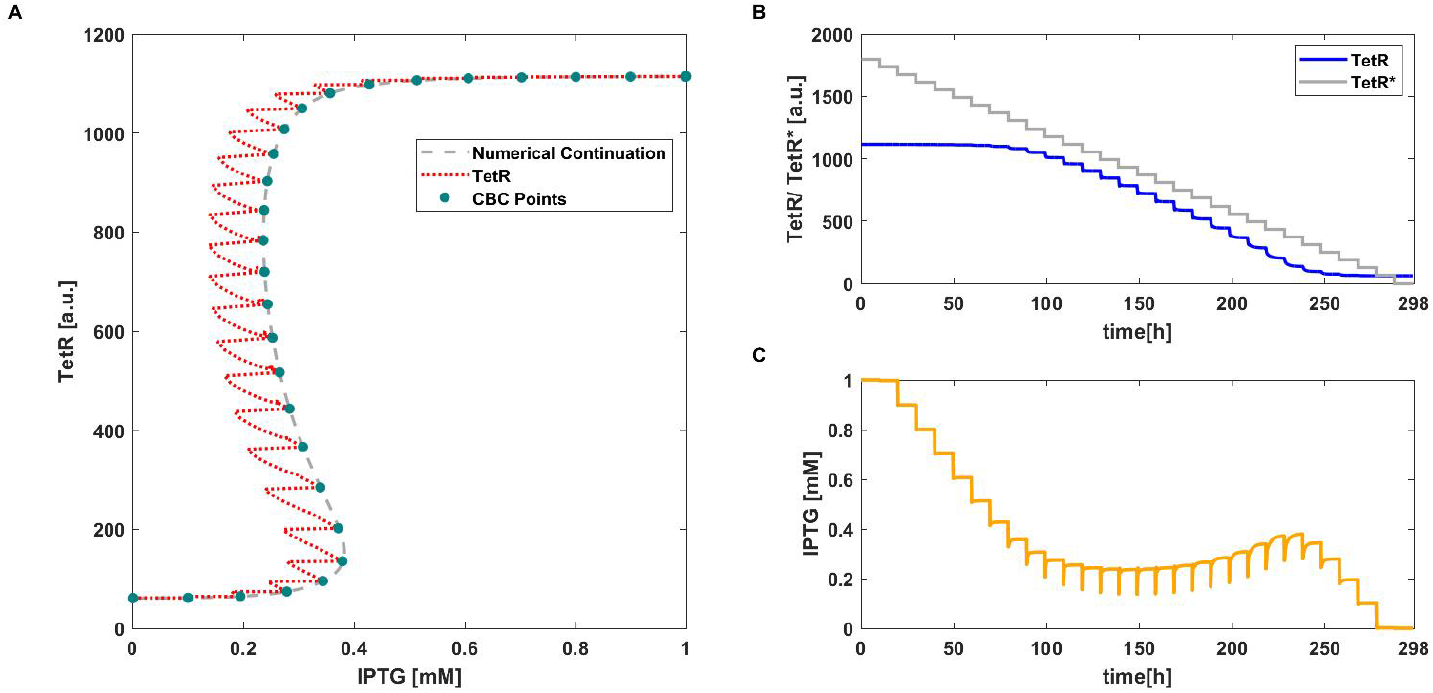
CBC with P controller applied to the deterministic toggle switch model (2). A) Equilibrium curve measured using CBC (·). *TetR*(*IPTG*) transient trajectories (..). Reference equilibrium curve obtained using numerical continuation (- -). B) Time evolution of *TetR* and the control reference signal *TetR\** (–). C) Time evolution of *IPTG* (i.e. control signal). Parameter values: *k_p_* = 0.0016 and *aTc* = 25*ngmL*^−^1.

**Figure 3:**
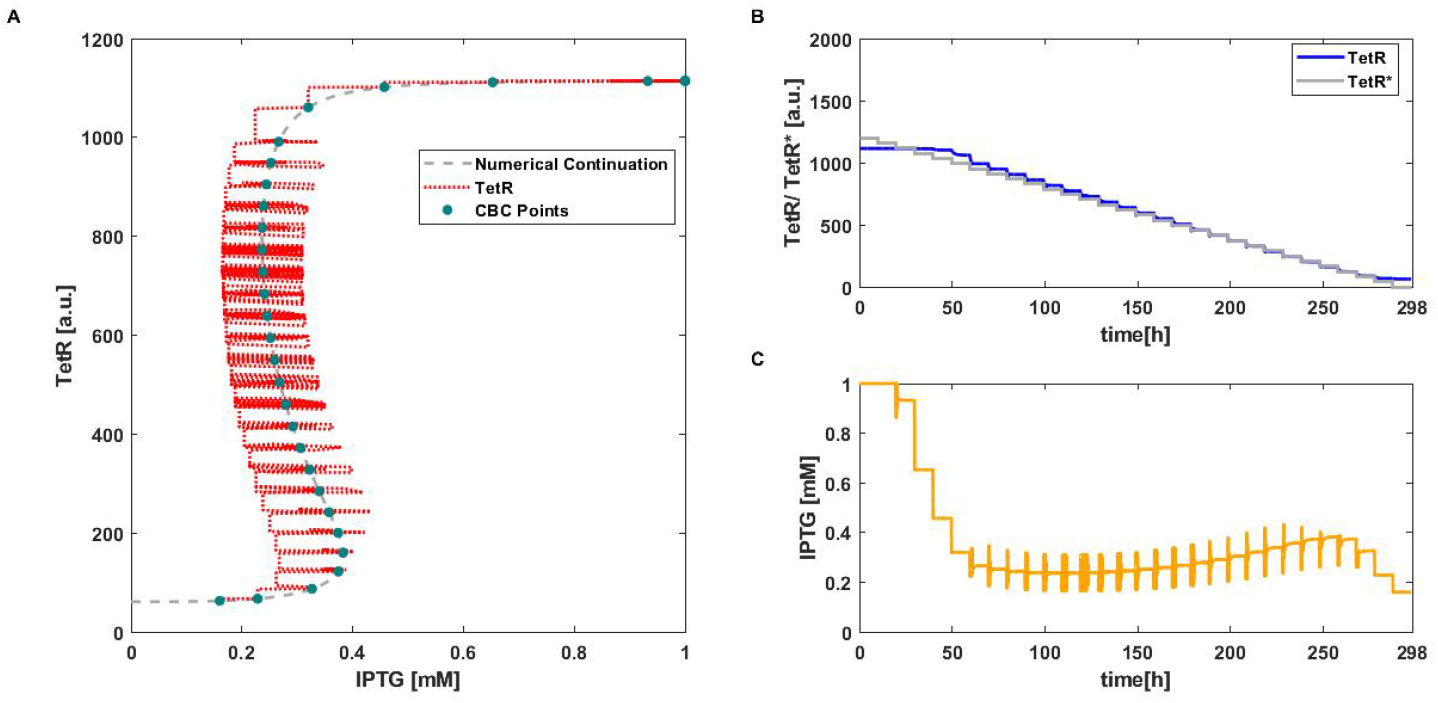
CBC with MPC applied to the deterministic toggle switch model (2). A) Equilibrium curve measured using CBC (·). *TetR*(*IPTG*) transient trajectories (..). Reference equilibrium curve obtained using numerical continuation (- -). B) Time evolution of *TetR* and the control reference signal *TetR\** (–). C) Time evolution of *IPTG* (i.e. control signal). Parameter values: *γ* = 0.3 (Eq. (25)) and *aTc* = 25*ngmL*^−^1.

The proportional control signal is linearly dependent on the error signal. Every time the reference is modified (Fig. 2 B), the error changes and consequently the control signal as well changes (Fig. 2 C). As the controller does not include an integral action, the error never nullifies, but it becomes constant once the system reaches the equilibrium. When this happens, the control signal and bifurcation parameter *IPTG* becomes constant as well (because the proportional contribution is constant), and the steady state can be collected.

With MPC, the control action is computed as the optimal signal that minimizes the error, and therefore its contribution pushes the system’s trajectories towards the reference signal, generating some early oscillations (Fig. 3 B, C). As the bifurcation parameter is not directly proportional to the control error, the latter does not have to be different from zero. The steady state error introduced by the MPC algorithm is much lower than the proportional one (see Fig. 2 B and Fig. 3 B); thus, it is easier to define the range of reference values for the MPC as it corresponds to the effective dynamical range explored by the continuation algorithm. Instead, when using a proportional controller, the range of the control reference may need to be varied significantly (as in Fig. 2 B, where the maximum reference was set to *TetR*\* = 1800[*a.u*.] in order for the algorithm to get a full coverage of the bifurcation diagram). This should not be considered as a fault of the proportional algorithm but only as a difference between the two control strategies. Results in Fig. 2 and Fig. 3 are also represented in Supplementary Movies S1 and S2.

To decrease the total time of each simulation we also implemented a steady state detection routine that changes the reference each time the system reaches steady state (Methods). This, together with a smaller number of collected points, allowed to decrease the experimental time to approximately 55 hours for both the controllers, see Fig. S1 and S2.

### CBC can reconstruct the toggle switch bifurcation diagram under a stochastic simulation setting

CBC is now demonstrated on a stochastic toggle switch model to reproduce conditions more similar to a real experiment. As noise can strongly modify results of a single continuation experiment, we consider 10 repetitions of the same experiment. In each we collect 30 points, with the steady states values computed as described for the ODE case.

The resulting plot is a density map made of all the collected steady states 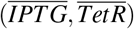 over the 10 simulations, that we can compare with the numerically computed bifurcation diagram (Fig. 4 A and 5 A). The results show dense clouds of points falling in the area where the branch of equilibria is, confirming the ability of the strategy to work also in a stochastic setting: both controllers were able to track the unstable branch of equilibria, preventing the system from jumping between the two stable steady states. For simplicity, we only show one out of 10 computed trajectories of *TetR* and *IPTG* in Fig. 4 B, C and 5 B, C.

**Figure 4:**
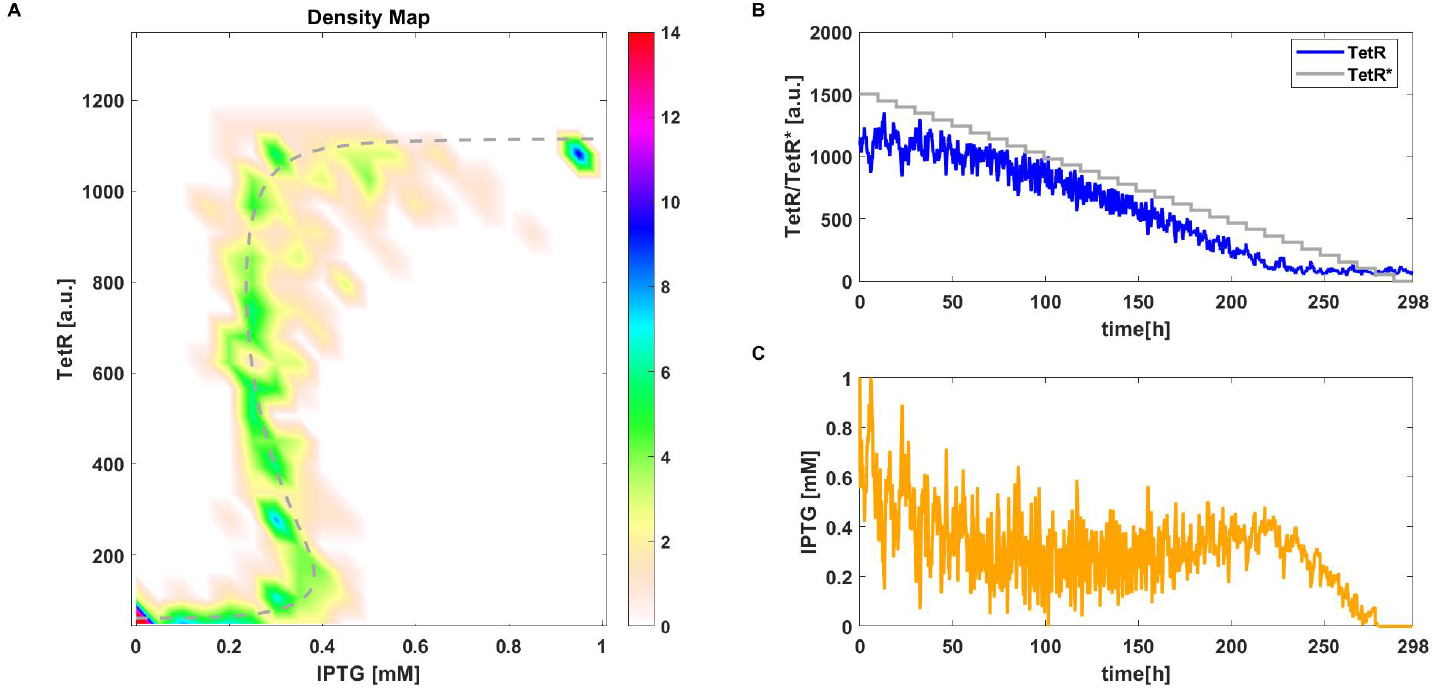
CBC with P controller applied to the stochastic toggle switch model (4). A) Density plot of equilibrium curve measured using CBC. Reference equilibrium curve obtained using numerical continuation (- -). B) Time evolution of one simulation of *TetR* and the control reference signal *TetR\** (–). C) Time evolution of *IPTG* (i.e. control signal). Parameter values: *k_p_* = 0.0016 and *aTc* = 25*ngmL*^−^1.

**Figure 5:**
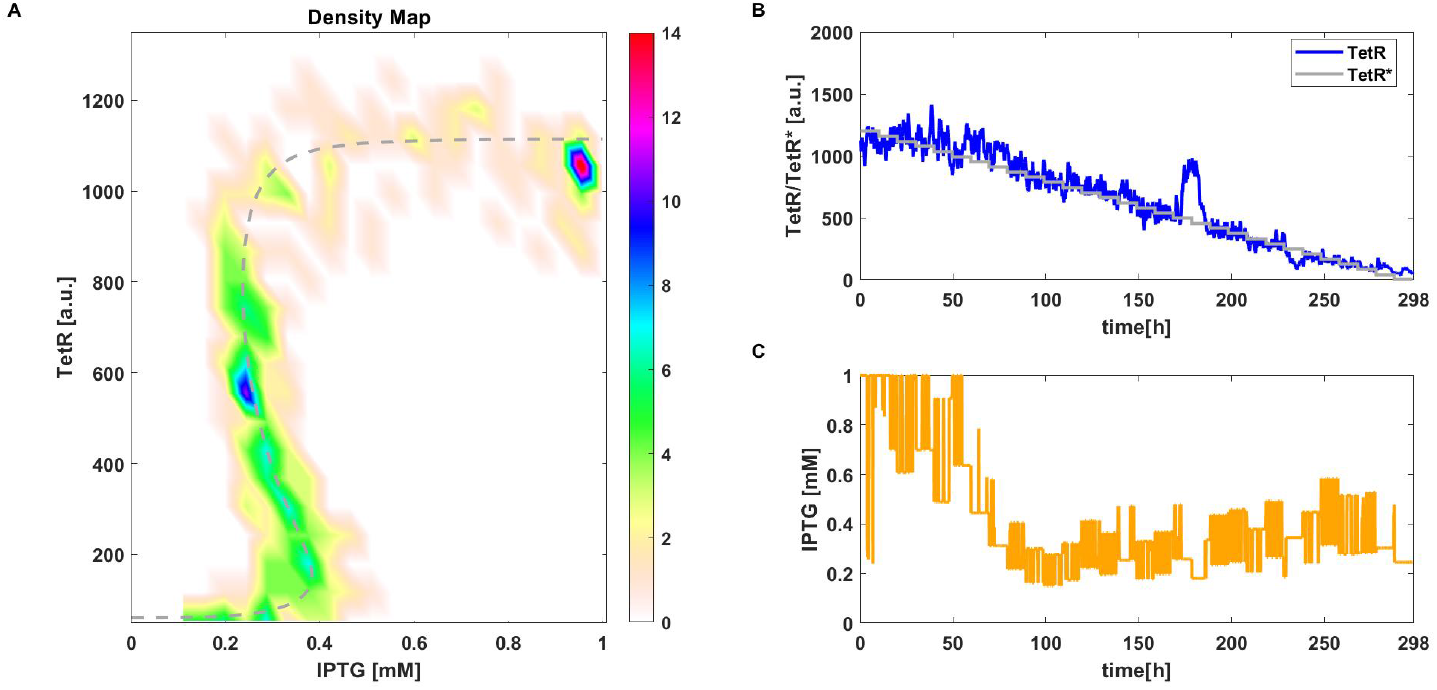
CBC with MPC applied to the stochastic toggle switch model (4). A) Density plot of equilibrium curve measured using CBC. Reference equilibrium curve obtained using numerical continuation (- -). B) Time evolution of one simulation of *TetR* and the control reference signal *TetR\** (–). C) Time evolution of *IPTG* (i.e. control signal). Parameter values: *γ* = 0.3 (Eq. (25)) and *aTc* = 25*ngmL*^−^1.

For both the proportional controller and MPC, the output signal shows oscillations due to noise, that also affect the control signal *u*. However, these oscillations do not seem to strongly alter the average values we take as steady state points. Similar observations to the deterministic case can be made. For the proportional controller, the error does not nullify and the reference signal is chosen in a wider range of values than for the MPC in order to uncover the full bifurcation curve (Fig. 4 B).

As in the deterministic scenario, the simulation time can be drastically decreased considering a variable step time associated to the computation of the steady states and a reduced amount of collected points. The resulting bifurcation curves can be seen in Fig. S3 A and S4 A, while panels B and C show just one representative trajectory of *TetR* and *IPTG*, respectively. Furthermore, for the stochastic scenario, we also considered the case of steady state check with full amount of points (30). We found that the reference shift guided by steady state detection could only reduce the total time to 228 hours for a proportional controller and 233 hours in case of MPC algorithm (see Figure S5 and S6), i.e. a 23.4% and a 21.8% reduction, respectively. Reducing the total testing time requires to decrease the number of points collected, which can be achieved without losing accuracy in the diagram retrieved as shown in Figs. S3 and S4. Nonetheless, the present method could be improved by developing a algorithm for automatic reference stepping with variable numbers of point collected.

### Single-cell CBC using agent-based simulations

To further prove the applicability of CBC to synthetic biology, we validated the method in BSim, a Java-based bacteria simulator [30, 33], where it is possible to consider a scenario more representative of a real experiment, including spatial distribution, movement, growth and division of cells as well as spatial distribution and diffusion of chemicals. We implemented single cell control considering the same model used for the stochastic scenario (Equation (4)). Simulation settings (i.e. sampling time, number of collected points, gains, etc.) where kept the same as above, with minor differences due to changes in the programming language.

In figures 6 B, 7 B and Supplementary Movies S3 and S4 representative examples of *TetR* and *IPTG* trajectories are shown. The bifurcation curves that are obtained, shown in Fig. 6 A and 7 A, are comparable with previous results obtained on the stochastic model. However, we observed worse control performances for the proportional controller in BSim. This is mainly due to additional factors introduced in simulations. Specifically, the cells biomechanics and their flush out from the microfluidic chamber are simulated, as well as the chemicals’ spatial distribution and diffusion that introduce additional delay in the control input delivery. Additionally, in BSim we explicitly simulated an actuation delay due to the time the media takes to be delivered to the cells within the microfluidic device. All these factors might contribute to the performances exhibited by the proportional controller. A similar result when using more realistic simulation environments was also found in [34], where the authors showed that the performance of a PI controller considerably deteriorated when the algorithm was tested on an agent based model, while an MPC algorithm was able to maintain similar performance. Further information about BSim can be found in the Methods section.

**Figure 6:**
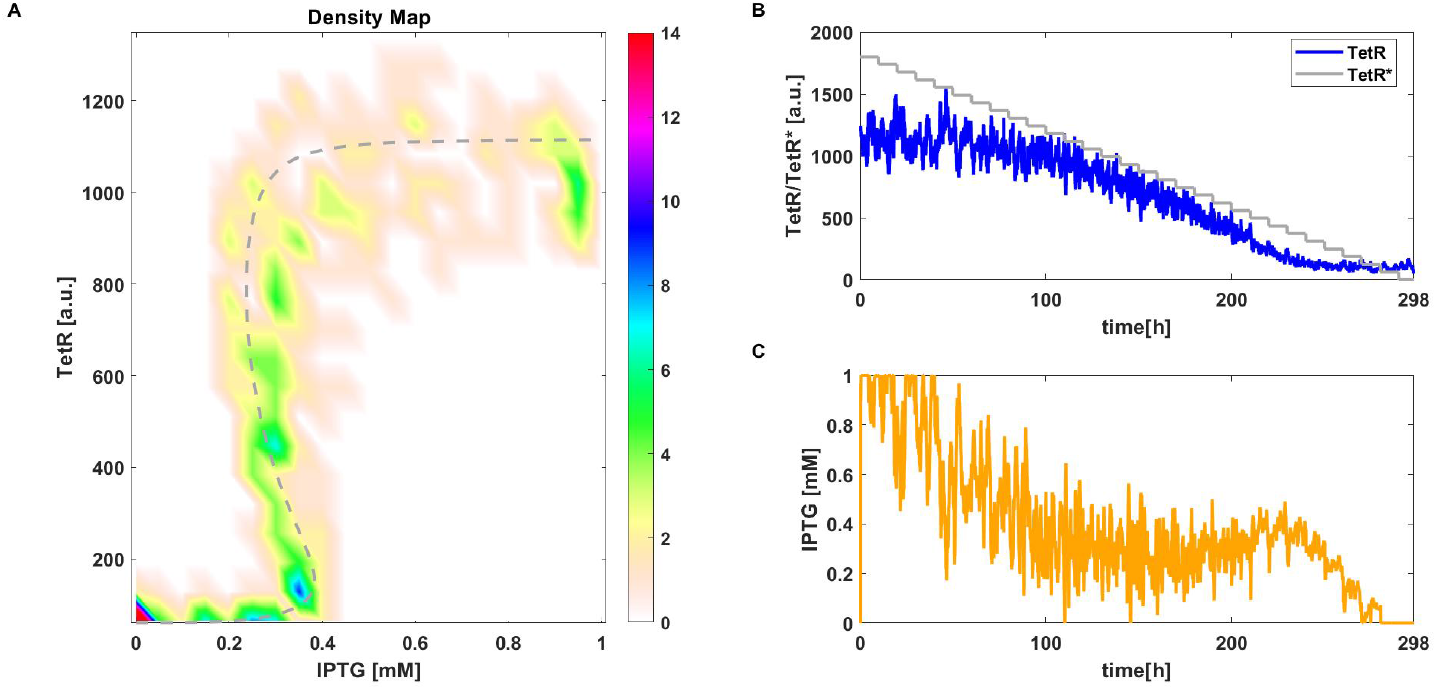
CBC with P controller applied to the stochastic toggle switch model (4) in BSim. A) Density plot of equilibrium curve measured using CBC. Reference equilibrium curve obtained using numerical continuation (- -). B) Time evolution of one simulation of *TetR* and the control reference signal *TetR\** (–). C) Time evolution of *IPTG* (i.e. control signal). Parameter values: *k_p_* = 0.0016 and *aTc* = 25*ngmL*^−^1.

**Figure 7:**
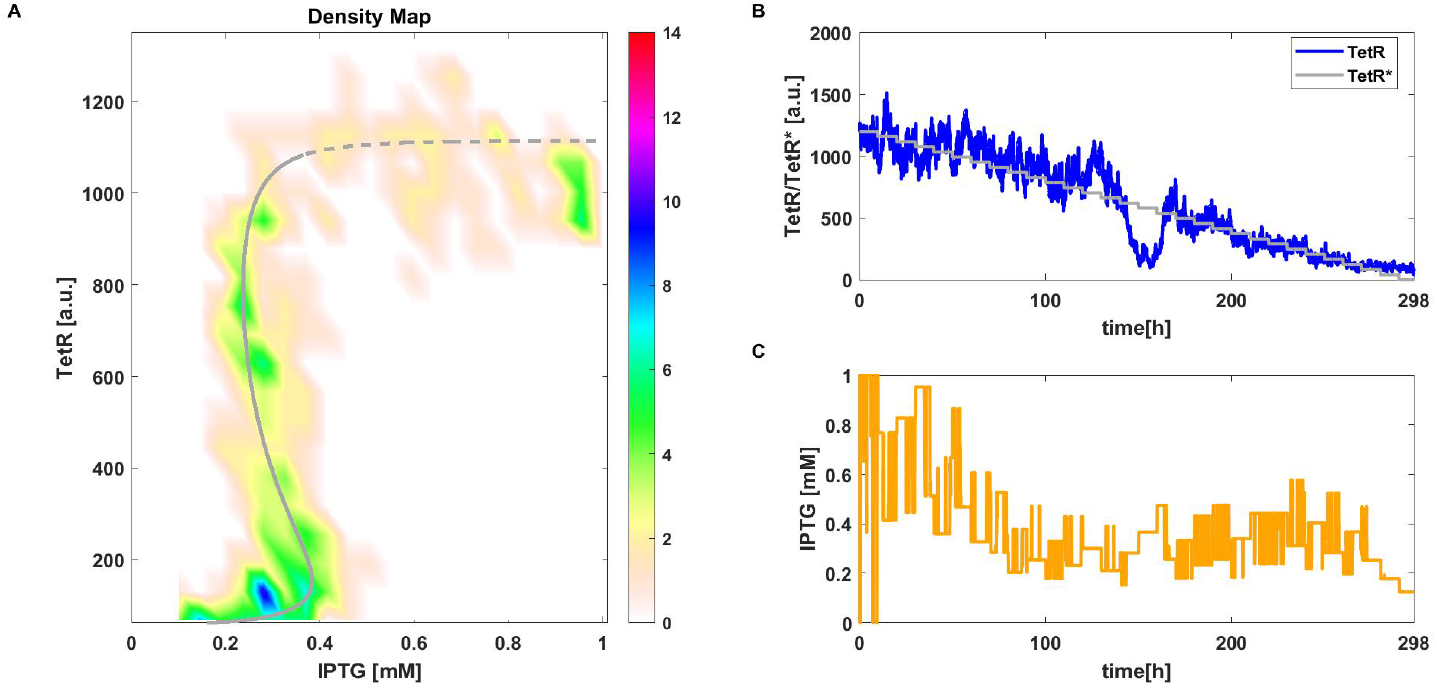
CBC with MPC applied to the stochastic toggle switch model (4) in BSim. A) Density plot of equilibrium curve measured using CBC. Reference equilibrium curve obtained using numerical continuation (- -). B) Time evolution of one simulation of *TetR* and the control reference signal *TetR\** (–). C) Time evolution of *IPTG* (i.e. control signal). Parameter values: *γ* = 0.3 (Eq. (25)) and *aTc* = 25*ngmL*^−^1.

Results with reduced points and steady state detection obtained in BSim (Fig. S7 and Fig. S8) were similar to short stochastic simulations in Matlab. Here, the increased complexity of simulations takes into account many sources of noise (such as spatial diffusion of chemicals); for this reason, results are more scattered around the numerically computed bifurcation diagram (see Fig. S7 A and Fig. S8 A). Nonetheless, implemented controllers manage to stabilise trajectories along the unstable branch of equilibria.

Considering the stochasticity of the experiments, there might be variability in the amount of time needed to uncover the bifurcation curve but, in general, it can be reduced up to less than a third of the original simulations.

### Model calibration via data collected using CBC in a noisy scenario

Methods for parameters identification in nonlinear systems often rely on optimization routines: given a model, parameters identification allows to calibrate its output in order to match experimental data (refer to [35, 36] for further information).

Using the approach by Beregi et al. [32], we show how the equilibria measured using CBC can be exploited to estimate the toggle switch model parameters, and how data corresponding to unstable equilibria helps retrieving a model that captures the bistable behaviour of the system. Further information about the identification process can be found in Methods.

The equilibrium curves obtained from parameters identified using CBC data are shown in Fig. S9 D for the deterministic scenario and in Fig. S9 E, F for the stochastic scenario (in green and red data associated to the proportional controller and to the MPC, respectively). For the sake of comparison, model parameters were also estimated from points collected using a traditional, open-loop parameter sweep where different levels of input parameter (IPTG) were tested for both noise-free and noisy scenarios (Fig. S9 A and B, C, respectively). The equilibrium curve obtained in this way generally does not reproduce the bistable behaviour of the system, except slightly in the ideal case of the deterministic simulations (Fig. S9 A). On the other hand, data collected with CBC still capture bistability even in stochastic simulations (Fig. S9 E, F), although without being able to reproduce the original numerical bifurcation curve.

Parameter estimation results show that the use of data including unstable equilibria helps estimating parameters that reproduce bistability. As such, it is thought that the MPC control approach is preferable compared to the proportional control as it offers greater control over the distribution of data points along the equilibrium curve. Estimated parameters are shown in Table S1 for all different collected data. Interestingly, the variation of some parameters does not significantly affect the resulting fitted curve. Take, for example, the red line in S9 D, which is in excellent agreement with the reference equilibrium curve but was calculated with some identified parameters values being up to 41% different from their nominal value, hinting to non-identifiability of some parameters. We confirmed this by performing a structural identifiability analysis on the system, using the STRIKE-GOLDD toolbox [37], which showed unidentifiability of 9 parameters out of 14 when we considered two measurable outputs (*LacI* and *TetR)*, one known control input (IPTG) and one other input as a fixed constant (aTc). For further reference, structural unidentifiability of the 2 ODEs toggle switch system for fixed inputs was also highlighted by Villaverde et al. [38].

We note that bifurcation curves estimated from sweeps can be improved to reproduce bistability if additional knowledge about the presence of horizontal asymptotes for the stable branches is introduced in the cost function used for parameter estimation (Fig. S10 A-C). However, obtaining qualitatively different results for similar estimation approaches would typically lead to poor confidence in the results. This modified approach is also not systematic as the true behaviour of the system is assumed to be known. Using this new approach with CBC data further improves the agreement between the reference and identified models (Fig. S10 D-F).

### Concluding Remarks

This paper presents the first application, *in silico*, of CBC to biochemical systems. In particular, CBC can track the stable and unstable equilibria of a synthetic gene network behaving as a toggle switch. In the absence of noise, a perfect agreement between equilibrium curves measured using CBC and standard numerical continuation methods is found. A notable challenge in biological applications, as compared to the mechanical systems studied so far with CBC, is the significant presence of (process) noise. Our results on the stochastic models (Matlab and Bsim) clearly show that CBC is still able to uncover the bistable nature of the system, as recorded data points qualitatively follow the equilibrium curve of the underlying deterministic model.

Besides exploring the nonlinear dynamics of the physical system directly in the experiment, CBC provides informative data for parameter estimation. Our results show that the measured data, especially those falling in the unstable region, was critical for the identification of model parameters that can reproduce bistability.

CBC appears therefore as a valuable tool for uncovering the dynamics of potentially more complex synthetic gene networks even in the presence of significant levels of noise. CBC can also be used to characterise oscillations [39] and other bifurcations [17, 40]. The information gathered with CBC has the potential to enable a more accurate understanding of biochemical interactions and thus a more precise prototyping of those behaviours into novel synthetic gene circuits.

An important feature of CBC is that it is a model-free method since it does not require a model of the system to work, and the accuracy of the results is independent from any modelling assumption. CBC also does not rely on a particular control strategy, provided the controller can be made noninvasive and is able to stabilise unstable responses. In this regard, the use of a mathematical model (even approximate) can improve the control performance. For instance, the initial estimate of a proportional controller gain can be obtained based on a model and then further refined by trial and error during experiments. We also showed that a model-predictive controller, based on a simple linear model, can provide a smaller steady-state control error, resulting in better discretisation of the equilibrium curve and beneficial for parameter estimation. MPC is a controller broadly used in biology [20] which generally performs best for fast varying references [41]. We hope that the use of MPC in the present paper paves the way for a wide range of applications of CBC in systems and synthetic biology.

### Future developments

In future studies we aim to apply the strategy *in-vivo*, using the microfluidic experimental set-up already described in [20, 22, 24], which proved to be successful for control experiments involving both mammalian and bacterial cells.

Several challenges will have to be addressed to enable a wider uptake of CBC:

1. *Noise*. The presence of noise can make stabilization more difficult, as observed here and also highlighted by Lugagne et al. [6]. Moreover, the presence of noise affects the noninvasiveness associated to the control input. Indeed, noise introduces variations in the control signal which cannot be suppressed completely. While this appears to have a mild effect on the system considered here, this is arguably not the case in general. As of now, some noise-related effects can be compensated by considering signal processing techniques, but more rigorous methods are required.
2. *Control*. As of now, there are no general and systematic methods to design the controller used in CBC. Currently, control parameters are found by trials and errors. While this works for simple systems, this approach does not scale to more complex systems where more sophisticated control strategies including, for instance, multiple inputs and outputs, will be needed. A step in that direction has recently been made by Li and Dankowicz [42] who proposed adaptive control design strategies for CBC.
3. *Experimental time*. Our initial simulations were particularly long (298 hours in Fig. 4-7). While such experiments can be achieved with some experimental set-ups – Balagaddé et al. [43] performed experiments on an *Escherichia coli* prey-predator system that lasted between 200 and 500 hours using a microchemostat platform – this is currently not possible using microscopy/microfluidic platform [6, 22]. It is in principle possible to significantly reduce testing time by improving the performance of the controller to reduce the transient. However, this can prove difficult due to the significant presence of noise in biochemical systems. Nonetheless, we showed that, for the presented gene network, it is possible to adjust the number of recorded points and CBC parameters to significantly reduce experimental time (2-3 days in the deterministic case, and 3-4 days in the stochastic case) while preserving a fine discretisation of the equilibrium curve.

## Methods

### Model of the toggle switch

The genetic toggle switch is a well-characterized bistable biological circuit, first embedded in *Escherichia coli* by Gardner et al. [10]. The system consists of two repressors (*LacI* and *TetR*), which mutually repress each other, and two inducers (Atc and *IPTG*), which can externally tune the genes’ production (Fig. 1 A). In the absence of inducers, the circuit exhibits two stable equilibria. Levels of *LacI* and *TetR* can be measured via fluorescent reporters: for example, *mKate2*(*RFP*) and *mEGFP*(GFP) can be used to monitor *LacI* and *TetR*, respectively [6].

Here, we will refer to the Hill-type model developed by Lugagne et al. [6], representing mRNA transcription, translation and degradation/dilution due to growth. Mathematically, the system is described by a set of four ordinary differential equations:

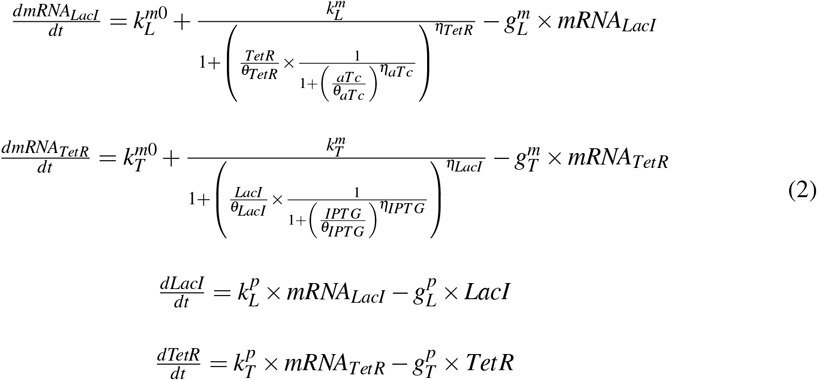

where the system states are given by the genes production *LacI* and *TetR* and the associated *mRNAs* (*mRNA_LacI_, mRNA_TetR_*), 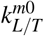 is the leakage transcription rate, 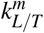 is the transcription rate, 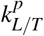 the translation rate, 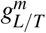 the *mRNA* degradation rate and 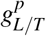 is the protein degradation rate. Parameters are listed in table 1.

**Table 1:** Parameters for the toggle switch model

| Parameters | Description | Value |
| --- | --- | --- |
| $k_L^{m0}$ | Transcription rates ($mRNA \min^{-1}$) | $3.20e - 2$ |
| $k_T^{m0}$ | - | $1.19e - 1$ |
| $k_L^m$ | - | 8.30 |
| $k_T^m$ | - | 2.06 |
| $k_L^p$ | Translation rates ($a.u. \ mRNA^{-1} \min^{-1}$) | $9.726e - 1$ |
| $k_T^p$ | - | 1.170 |
| $g_L^m$ | Degradation rates ($\min^{-1}$) | $1.386e - 1$ |
| $g_T^m$ | - | $1.386e - 1$ |
| $g_L^p$ | - | $1.65e - 2$ |
| $g_T^p$ | - | $1.65e - 2$ |
| $\theta_{LacI}$ | $plac$ regulation by TetR (-) | 31.94 |
| $\eta_{LacI}$ | - | 2.00 |
| $\theta_{IPTG}$ | - | $9.06e - 2$ |
| $\eta_{IPTG}$ | - | 2.00 |
| $\theta_{TetR}$ | $ptet$ regulation by LacI (-) | 30.00 |
| $\eta_{TetR}$ | - | 2.00 |
| $\theta_{aTc}$ | - | 11.65 |
| $\eta_{aTc}$ | - | 2.00 |
| $k_{IPTG}^{in}$ | IPTG exchange rate ($\min^{-1}$) | $2.75e - 2$ |
| $k_{IPTG}^{out}$ | - | $1.11e - 1$ |
| $k_{aTc}^{in}$ | aTc exchange rate ($\min^{-1}$) | $1.62e - 1$ |
| $k_{aTc}^{out}$ | - | $2.00e - 2$ |

The authors assumed that the repressors could be modelled with Hill function (*h*(*x, θ, η*) = 1/(1 + *x*/*θ*)^*η*^), where *θ* represents a threshold parameter for the Hill function and η is the Hill coefficient. Two more equations were added to take into account inducers diffusion through the cell membrane. Formally:

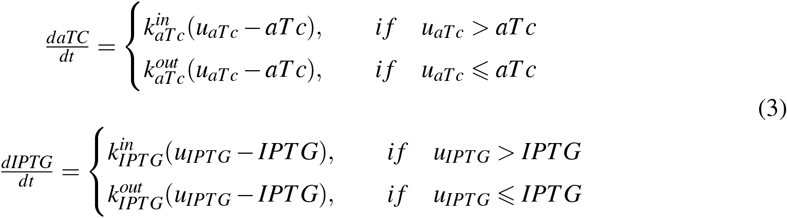

The deterministic version of the model does not capture the intrinsic stochasticity of biochemical processes. A more accurate description is provided by the stochastic modelling procedure, based on the pseudo-reactions listed in table 2.

**Table 2:** Table of pseudo-reactions: here 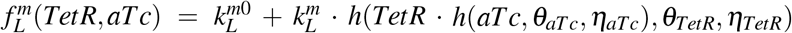 and 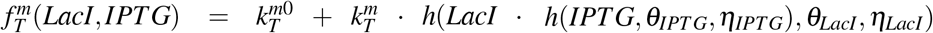 are gene regulation function, and *h*(*x, θ, η*) = 1/(1 + *x*/*θ*)^*η*^ is the decreasing Hill function. The others arrows superscripts are parameters that can be found in Table 1.

| Description | Pseudo-reaction |
| --- | --- |
| transcription | $\emptyset \xrightarrow{f_L^m(TetR, aTc)} mRNA_L$<br>$\emptyset \xrightarrow{f_T^m(LacI, IPTG)} mRNA_T$ |
| translation | $mRNA_L \xrightarrow{k_L^p} mRNA_L + LacI$<br>$mRNA_T \xrightarrow{k_T^p} mRNA_T + TetR$ |
| degradation/dilution | $mRNA_L \xrightarrow{g_L^m} \emptyset$<br>$mRNA_T \xrightarrow{g_T^m} \emptyset$<br>$LacI \xrightarrow{g_L^p} \emptyset$<br>$TetR \xrightarrow{g_T^p} \emptyset$ |

Specifically, we employed a SDE based model for the description of biochemical systems [44]:

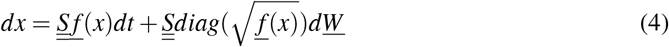

Here:

- 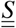 is the stoichiometry matrix where each row is associated to a state variable and each column *i* includes stoichiometry coefficients associated to the reaction *i*.
- 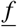 is a vector where each element *i* corresponds to a propensity function. The latter is a function describing the probability of a certain reaction given the concentration of the chemical species ([45]).
- 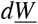 is a vector containing the Wiener Process increments.

Pseudo-reactions and propensity functions were all given in [6], and here reported in Table 2. All simulations were performed using Matlab R2019b. To solve the system (2) we used the function ode45, while for the stochastic model in eq. (4) we implemented the Euler-Maruyama method [44], that is known to be slightly less accurate than the Stochastic Simulation Algorithm [45], but more computationally efficient.

Throughout the paper we have assumed *aTc* to be fixed to 25*ngmL*^−^1 for all simulations.

### Control-based continuation theory: using control to track equilibria of the underlying uncontrolled system

CBC seeks to define a bifurcation diagram by the application of a noninvasive control action, that does not modify the underlying uncontrolled system’s equilibria position in parameter space. To do so, the original method looks for a control action whose contribution vanishes asymptotically (eq. (1)). However, the same noninvasiveness can be achieved with a slightly simplified method, provided that the control action enters the system as bifurcation parameter [31]. The fundamental principles of the simplified CBC method used in this paper can be explained using the scalar differential equation:

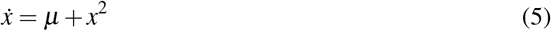

where *μ* ∈ ℝ is the bifurcation parameter. Eq. (5) corresponds to the normal form of a fold (or saddle-node) bifurcation. The equilibria of the above system are given by the formula:

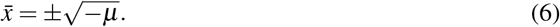

To trace out the equilibrium curve of (5), including both stable and unstable equilibria, CBC relies on the presence of a stabilising feedback controller. The equation of motion of the system including feedback control is given by:

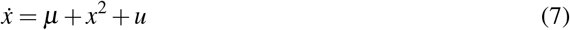

where *u* is the control signal given by a suitable, i.e. stabilizing, control law. When the control signal and bifurcation parameter affect the system in the same way (as in Eq. (7)), the static component of the control signal can be interpreted as a shift of the bifurcation parameter *μ*. In this paper, a simple linear proportional (P) law will be considered such that *u* is given by

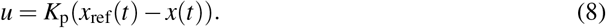

where *x*_ref_(*t*) is the control reference (or target) and *K_p_* is the proportional gain. Using (8) as control input, (7) becomes:

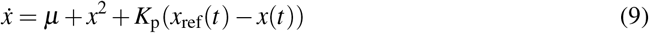

that can be reordered as:

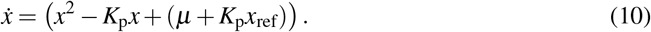

Thus, we can write the equilibria of (10) and the control signal at steady state as:

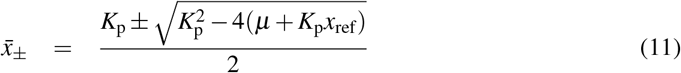

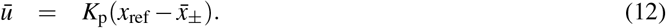

Writing *x*_ref_ as function of the steady state control action u, we can substitute Eq. (12) in Eq. (11) to obtain:

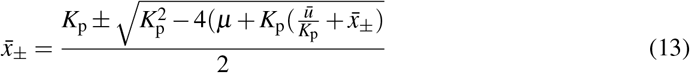

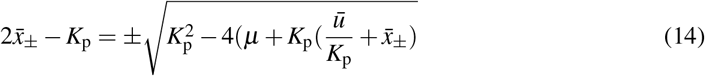

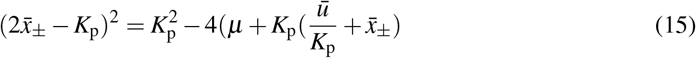

Solving the squared term and simplifying the equation we can thus obtain:

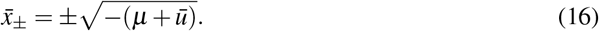

Eq. (16) defines the same equilibrium curve as Eq. (6) for a parameter 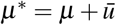. Therefore, the steady state contribution of the control signal does not have to be removed and it can simply be considered as a shift in parameter space. The control strategy so defined is noninvasive (it does not modify the position of system’s equilibria in parameter space) and the requirement of eq. (1) can be translated into:

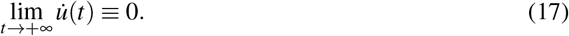

which is equivalent to converging to a steady state value.

Here we demonstrated how CBC can be applied to track equilibria of a system by the application of a stabilizing proportional controller. However, the same results could be achieved with other control strategies as long as the controller stabilises the system’s unstable equilibria. In fact, CBC is not restricted to a particular form of control and thus the control law can in principle take a general form:

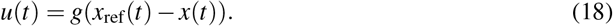

### Main steps of the control-based continuation algorithm

The algorithm applied for control-based continuation can be briefly summarized in the following steps:

1. Set a new control reference. A reference value *x*_ref_ = *TetR\** is used to evaluate the error (the difference between the measured *TetR* and the reference target), and it is given to the controller. The range where the reference is picked normally covers the minimum and maximum expression of the protein of interest, but it can be modified according to the need (Fig. 2 B shows a maximum reference value of *TetR*\* = 1800[*a.u*], although the maximum expression of *TetR* is 1200[*a.u*]).
2. Compute the control action. Depending on the control strategy, either the MPC performs optimization and defines the optimal control input or the proportional controller evaluates the control action depending on the error value.
3. Feed the control action to the system. The computed control action *ū* is fed to the system for 5 minutes continuously, as it is considered to be the minimum sampling time for measurements and actuation in an hypothetical microfluidics/microscopy experimental set-up. After 5 minutes, the system output is measured and a new error is computed. Steps 2 and 3 are repeated until the process is considered at steady state.
4. Check for steady state. The initial implementation of the algorithm considers the system at steady state after a fixed amount of time (9*h* and 55*min* for experiments shown in the Results section). The algorithm allows a variable time for steady state computation. In this case, to compute the steady state of *TetR* and *IPTG*, the slope of the linear curve fitting the last 12 samples after 3 hours of simulation is calculated for both the error and the control signals (where the slope corresponds to the angle the curve makes with respect to the *x* – *axis*). If the computed value is below a user-defined tolerance, the system is assumed at steady state. Otherwise, the simulation continues and the steady state is checked every new sample, until the slope is sufficiently small. A threshold is set on both the error and the control slopes, which may vary between a deterministic and a stochastic simulation.
5. Acquire the steady state. The steady state values of *TetR* and *IPTG* (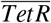 and 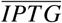) are saved as the average value computed over the last 12 samples of each signal, and the algorithm goes back to step 1.

All steps are repeated until enough steady states are collected, or until the range of output and parameter values has been covered. An initial guess is given as initial state for the algorithm to start with. A schematic of few steps can be seen in Fig. 1 B.

Results of the variable stepping reference algorithm can be seen in Fig. S1-S8 for both the deterministic and the stochastic scenarios.

### Data availability

The code used in this publication is available at: https://github.com/lrenson/cbc-synbio-paper, https://github.com/diBernardoGroup/Control-Based-Continuation-of-a-genetic-Toggle-Switch. Movies S1, S2, S3 and S4 are provided as supplementary files.

### Proportional Controller

The simplest controller that can satisfy CBC requirements is a proportional controller, formally described by the control law of Equation (8). The error signal is computed by subtracting the measured output to the control reference (*TetR*\* – *TetR*). The error is then fed to the controller to evaluate the control input *IPTG* given to the toggle switch system. A schematic of the control loop can be seen in Fig. S11 A.

Despite its simplicity, the proportional controller has proved to be a solid choice for CBC. As long as the controller can stabilise the system, the control action is noninvasive and thus the points collected through the CBC routine correspond to the equilibria of the underlying uncontrolled system.

To tune the proportional gain *K_p_* we linearized the system around an unstable equilibria and then we used the root locus to find the minimum gain able to stabilise the linearized system. Depending on the type of simulation, deterministic or stochastic, the *K_p_* gain is then adjusted by trial and error.

### Model Predictive Controller

Model predictive control is a control scheme based on two main features: prediction and optimization. At each step, a model reproducing the process behaviour is used to predict the process outputs to given input signals. The input minimizing a cost function is computed using an optimization algorithm and fed to the controlled process. We chose a linear model to reproduce behaviours we expect from the real process. A further requirement for the prediction procedure is the process current state which, if not fully measurable, can be estimated using a Kalman filter. A schematic of the control loop can be seen in Fig. S9 B.

### System identification to reproduce the process experimental behaviour through a simplified model

The MPC strategy requires a model to predict the process trajectories and carry out the optimization routine. We artificially reproduced the experimental behaviours using stochastic numerical simulations. Specifically, we measured the response of the system’s outputs (*LacI* and *TetR*) to different randomly generated pulses of *aTc* and *IPTG*, both with fixed and variable amplitude. This data-set was used to identify a LTI system via noniterative subspace estimation algorithm, using the Matlab System Identification toolbox. The result was a continuous-time identified state space model of the following form:

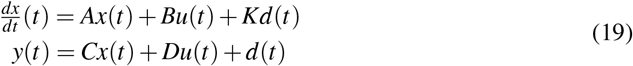

where *x*(*t*) is the state, *u*(*t*) is the input to the system, *y*(*t*) is the system output, *d*(*t*) is the state disturbance and *K* is its associated matrix [46].

Here, *A,B,C* and *K* are free parameters to estimate, while *D* was set to 0. Equation (19) is called innovation form of the state-space model and the matrix *K* corresponds to the Kalman gain matrix associated to the identified system.

### Definition of the Kalman predictor to reconstruct unmeasurable states

Considering the Toggle Switch system (2), we measure only *LacI* and *TetR* concentrations. To estimate the remaining states we opted for a Kalman filter, defined as:

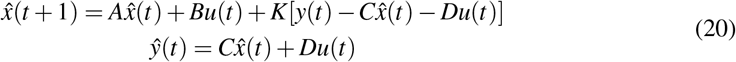

where 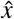 is the state estimate, 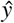 is the output estimate and all the matrices *A, B, C, D, K* correspond to those obtained through the identification process.

Eq. (20) can be rewritten as follows:

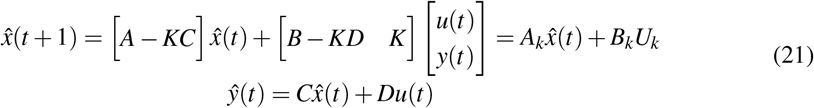

The addition of a correction based on the measured outputs *LacI* and *TetR* (*U_k_* = [*u*(*t*); [*LacI*, *TetR*]]) enables a better fit of the original process outputs [47].

The ability to predict the data recorded is evaluated using the percentage of the output variation that is reproduced by the model. This is defined as:

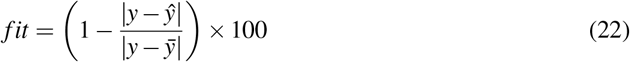

where *y* is the measured output, *ŷ* is the simulated predicted model output and 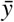 is the mean value of *y*. Fit percentages of *LacI* and *TetR* outputs for the system in eq. (21) are 96.4 and 92.2 respectively.

### Definition of a cost function to compute an optimal control action

To compute the optimal control action we employed a genetic algorithm [48]. Commonly, MPC algorithms use quadratic cost functions weighting both the state of the system and the control action. Considering a scalar example, a quadratic cost function could be defined as:

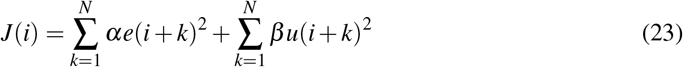

Here, *α* and *β* correspond to weighting coefficients reflecting the relative importance of the error *e* and the control input *u*; *N* represents the prediction horizon, and *i* the current time. To adapt the cost function to CBC and guarantee noninvasiveness of the control input we initially replaced the control input *u* by its variation Δ*u* as follows:

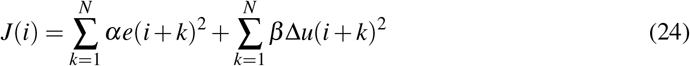

In this way, the optimization finds the best trade off between regulation accuracy and control variation. Although eq. (24) granted us good results in most of the simulations, the tuning of both *α* and *β* was difficult, as small changes into the CBC algorithm variables (such as the number of points collected during a single simulation) affected the quality of the results and demanded for retuning of the cost function parameters.

To simplify the cost function, we set *β* = 0 and *α* = *N* – *k* + 1 and forced the optimizer to limit both the control input amplitude and variation at each iteration point. Formally we defined the following constraints:

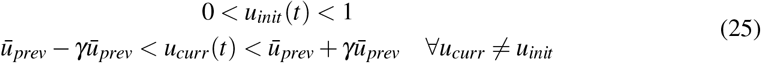

where *u_init_*(*t*) is the control input during the first step of the algorithm; *u_curr_*(*t*) is the control input for all other steps except for the first one and 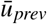 is the control input steady state value, saved before stepping the reference in the algorithm.

The presence of external constraints on the control action allows to have a noninvasive contribution without the need for the *β*Δ*u*(*i* + *k*)^2^ term in the cost function. The parameter *γ* defines a percentage bandwidth that we consider acceptable for the control action to vary between. In other words, at each iteration of the CBC algorithm -by which we mean every time we change the reference signal- the control action cannot vary more than *γ* with respect to the previous registered steady state input. In our simulations *γ* was set to 0.3 for 30 points (Fig. 3, 5) and to 0.5 for 11 and 12 points (Fig. S2, S4). Other settings for *γ* might be found in Figures captions. Fewer points result in larger control reference steps, which can be associated with bigger steps in the control input, and for this reason *γ* can be more easily calibrated just by considering the amount of points to be collected in a simulation.

### Parameters estimation process

For the estimation process, we calculated the analytical solutions of our toggle switch model (2) that will be used to build the cost function for parameter estimation. In particular, we want to define the function:

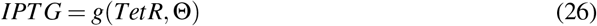

that has a unique real solution for each admissible value of *TetR* and for a chosen set of parameters **Θ** = (*klm*_0_, *klm*, *θ_aTc_*, *η_aTc_*, *θ_TetR_, η_TetR_, ktm*_0_, *ktm*, *θ_IPTG_, η_IPTG_, θ_LacI_, η_LacI_, klp, ktp*), which is the subject of the estimation process. The full Equation (26) is presented in the Supplementary (S1). Here we considered the parameters (*gtm, glm, glp, gtp*) to be fixed to the values computed in [6] and therefore we did not estimate them.

Once we computed the analytical definition of *g*(*TetR*, **Θ**), we built the following cost function:

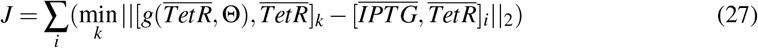

where 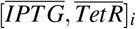 are the measured steady state values of control input and control output out of the CBC routine for the *i* point of the bifurcation curve. The *k* index looks for the estimated point 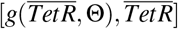 with minimum distance (L^2^ norm) from the measured steady state considered by the index *i*. 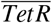 is normalized between 0 and 1 to be comparable with 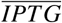.

Minimizing the cost function (27) for all points collected during an experiment, it is possible to estimate the best set of parameters **Θ** that allows to characterize a model of the toggle switch reproducing the bistability behaviour seen via experiments. Once a set of parameters is found (using a genetic algorithm), it is possible to build the bifurcation curve by solving Equation (26) for fixed values of *TetR*. Sets **Θ** that gave complex values of *IPTG* were directly discarded. Estimates of parameters for different experimental settings and percentage variation with respect to nominal values can be seen in Table S1.

In order to validate the results of the estimation using data collected with the CBC algorithm, a comparison with the parameter sweeps method was made. Parameter sweeps is a simple experimental design to collect data at different values of a parameter of interest and is commonly used for parameter estimation. Considering the example of the toggle switch, parameter sweeps is implemented registering the steady state signal 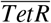 for fixed values of the input 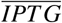. Collected points 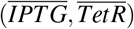 fall on the stable branches of our bifurcation curve, as shown in Fig. S11 A, B, C. This method does not allow to collect unstable steady states as there is no control over the system.

We showed that CBC grants a more robust estimation by collecting data in the unstable branch, which is not possible through simple parameter sweeps, and allows to retrieve the bistability of the system (see Fig. S9 D, E, F).

As highlighted in the main text, assuming an *a priori* knowledge of the system’s bistability, we could improve the performances of our calibration by releasing the constraint on sets **Θ** that generate complex values of *IPTG*. Furthermore, more complex cost function could be considered (for example, the minimum distance of measured point from the calibrated bifurcation curve). Fig. S10 shows this latter improved case, with very good fitting performances compared to the initial estimation. Although the noticeable improvement, the modified cost function requires a much longer computational time and knowledge of the system of interest’s dynamics, which is often not the case when CBC is applied experimentally.

### Agent-based simulations

All agent based simulations were performed using BSim, an agent-based environment designed to simulate bacterial populations [30, 33]. Here, cell bio-mechanics are explicitly modelled, together with cell growth and division. The spatial configuration and dynamics of the chemical inducers added in the culture environment are also simulated, introducing an extra time delay in the delivery of the inducer molecules to the cells. In addition, it is possible to mimic microfluidic-based experimental platforms, including related physical and technological constraints (e.g. dimensions and shape of microfluidic chambers, flush out of cells). Specifically, we can define the geometry and dimension of the microfluidic chamber, and the physical interactions of the cells with the device. We can account for cells movement and collision inside the growth chamber and simulate the flushing out of the cells from the chamber with their consequent removal from the set of analysable cells. The CBC algorithm was implemented by adding to the simulation environment both the proportional and the MPC controllers.

As previously done in Shannon et al. [22] and Salzano et al. [49], the simulation environment has been complemented with the SDE solver based on the Euler-Maruyama method (4).

BSim simulates experiments via a microfluidic-based system composed by a microfluidic device, a microscopy, a computer and an actuation system. Dimensions of the growth chamber were initially taken from Lugagne et al. [6] but then set to 16*μM* × 1.5*μM* × 1*μM* to reduce the simulation time. The proportional controller was implemented directly in Bsim, while the MPC was implemented in Matlab and externally called by the agent-based simulator. All bio-mechanical parameters of the cells were set to the same values used by Matyjaszkiewicz et al. [30]. Moreover, realistic constraints on the sampling and actuation time were considered, similarly to what already done for Matlab CBC simulations. More precisely, we enforced the state of the cells to be measured every 5min and we assumed the input provided to be changed at most every 5min. Finally, we assumed a 40s delay on the actuation of the control inputs, accounting for the time the inducers would take to reach the microfluidics cell culture chambers in a physical experiment.

All the factors taken into account in BSim added additional sources of variability to the simulations, sometimes affecting the performance of the control algorithms, as discussed in the Results and Discussion sections.

## Associated Content

Supporting Information. Figures of short CBC simulations in a deterministic setting (Fig. S1 and S2) and in a stochastic setting (Fig. S3 and S4) with both controllers and reduced points. Figures of short CBC simulations in a stochastic setting (Fig. S5 and S6) with both controllers and all points. Figures of short CBC simulations in a stochastic setting with BSIM (Fig. S7 and S8). Equilibrium curve computed for model parameters estimated with constraints on IPTG complexity using parameter sweep data, CBC with MPC data and CBC with P controller data (Fig. S9). Equilibrium curve computed for model parameters estimated without constraints on IPTG complexity using parameter sweep data, CBC with MPC data and CBC with P controller data (Fig. S10). Control Feedback Loops, both MPC and P, implemented for CBC (Fig. S11). Estimated parameters for the toggle switch model depending on the method used to collect data (Tab. S1). Full IPTG = *g*(*TetR*, **Θ**) equation for parameter estimation. CBC movies legends.

## Abbreviation

CBC: control-based continuation
MPC: model predctive control
P: proportional control

## Author Information

### Corresponding Authors

Ludovic Renson - *Department of Mechanical Engineering, Imperial College London, UK*. Co-senior author.

Lucia Marucci - *Engineering Mathematics Department, School of Cellular and Molecular Medicine and BrisSynBio, University of Bristol, UK*. Co-senior author.

### Authors

Irene de Cesare - *Engineering Mathematics Department, University of Bristol, UK*.

Davide Salzano - *Department of Electrical Engineering and Information Technologies, University of Naples Federico II, Italy*.

Mario di Bernardo - *Department of Electrical Engineering and Information Technologies, University of Naples Federico II, Italy*. Co-senior author.

### Author Contributions

I.d.C. designed and carried out the simulations in Matlab2019b and wrote the manuscript with inputs from all the authors. D.S. developed the BSim code and wrote the agent-based simulations section. M.d.B supervised the initial development of CBC and the agent-based implementation. L.R. provided the initial framework for the CBC code. L.R. and L.M. conceived the project and supervised the entire work.

## Acknowledgments

I.d.C. is supported by an EPSRC DTP Scholarship. L.M. is supported by the Medical Research Council grant MR/N021444/1, by the Engineering and Physical Sciences Research Council grants EP/R041695/1 and EP/S01876X/1, and by the European Union’s Horizon 2020 under Grant Agreement No. 766840. L.R. is funded by a Research Fellowship from the Royal Academy of Engineering (RF1516/15/11).

