## Supplementary Information for "Control-based continuation: a new approach to prototype synthetic gene networks"

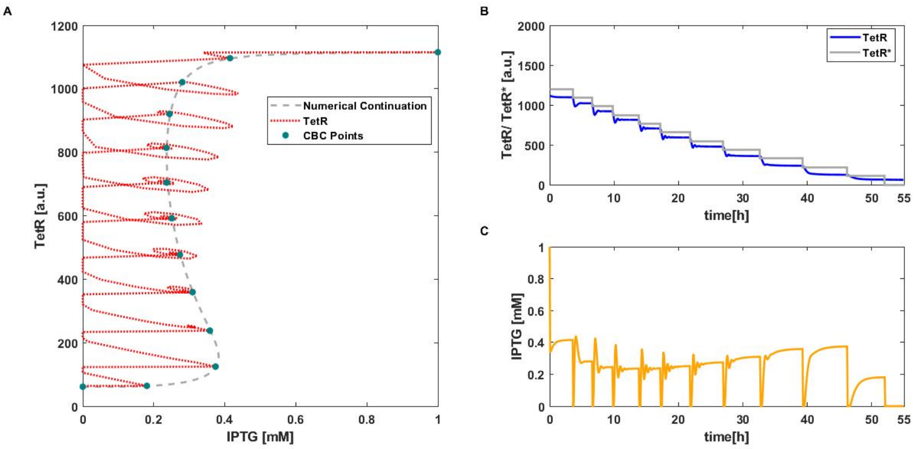


Figure S1: Short simulation of CBC with P controller applied to deterministic toggle switch model (2). A) Equilibrium curve measured using CBC (·). $TetR(IPTG)$ transient trajectories (..). Reference equilibrium curve obtained using numerical continuation (- -). B) Time evolution of $TetR$ and the control reference signal $TetR^*$ (–). C) Time evolution of $IPTG$ (i.e. control signal). Parameter values: $k_p = 0.004$ and $aTc = 25ngmL^-1$.

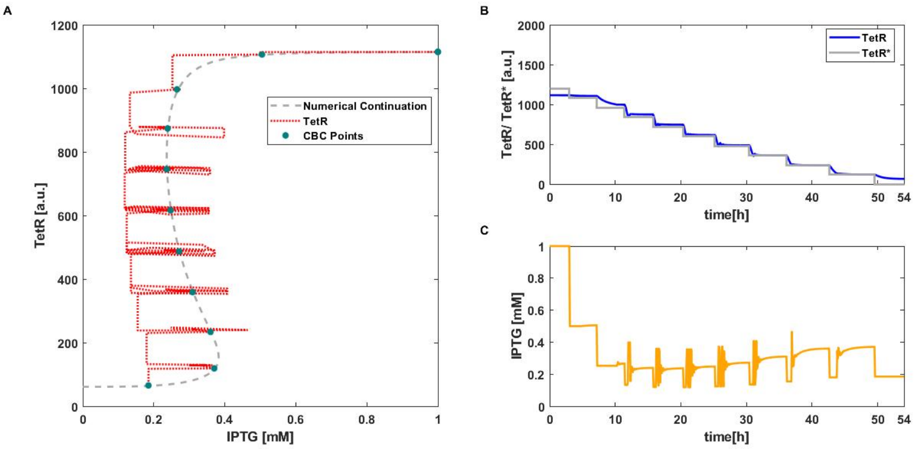


Figure S2: Short simulation of CBC with MPC applied to deterministic toggle switch model (2). A) Equilibrium curve measured using CBC (·). $TetR(IPTG)$ transient trajectories (..). Reference equilibrium curve obtained using numerical continuation (- -). B) Time evolution of $TetR$ and the control reference signal $TetR^*$ (–). C) Time evolution of $IPTG$ (i.e. control signal). Parameter values: $\gamma = 0.5$ (Eq. 25) and $aTc = 25ngmL^-1$.

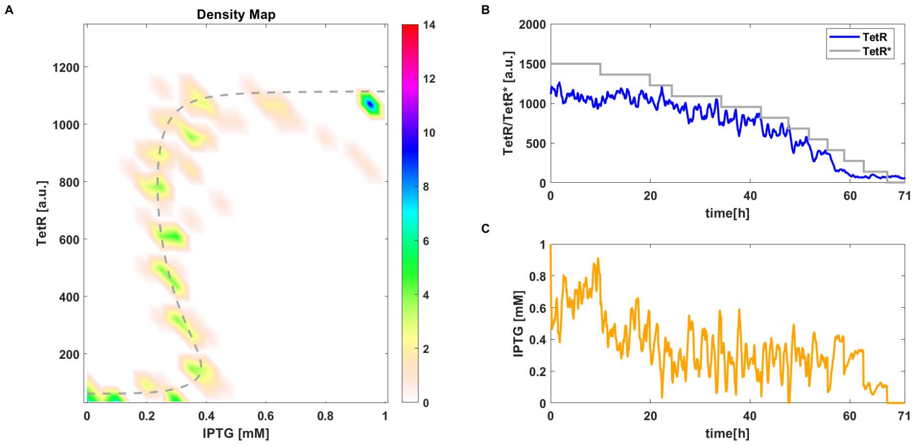


Figure S3: Short simulation of CBC with P controller applied to stochastic toggle switch model (4). A) Density plot of equilibrium curve measured using CBC (12 points each simulation). Reference equilibrium curve obtained using numerical continuation (- -). B) Time evolution of one simulation of $TetR$ and the control reference signal $TetR^*$ (–). C) Time evolution of $IPTG$ (i.e. control signal). Parameter values: $k_p = 0.0016$ and $aTc = 25ngmL^{-}1$.

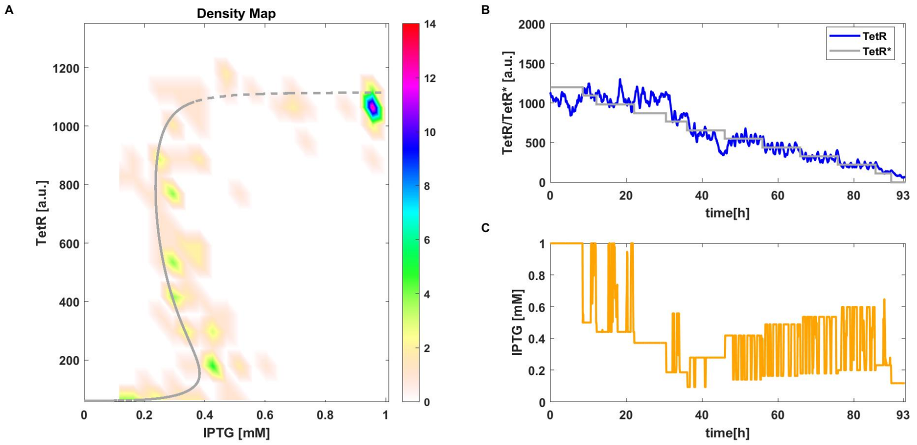


Figure S4: Short simulation of CBC with MPC applied to stochastic toggle switch model (4). A) Density plot of equilibrium curve measured using CBC (12 points each simulation). Reference equilibrium curve obtained using numerical continuation (- -). B) Time evolution of one simulation of $TetR$ and the control reference signal $TetR^*$ (–). C) Time evolution of $IPTG$ (i.e. control signal). Parameter values: $\gamma = 0.5$ (Eq. 25) and $aTc = 25ngmL^{-}1$.

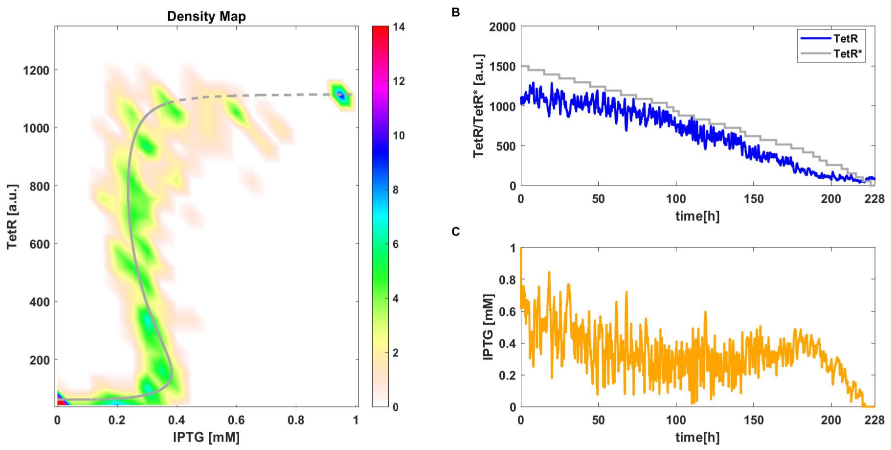


Figure S5: Short simulation (all points) of CBC with P controller applied to stochastic toggle switch model (4). A) Density plot of equilibrium curve measured using CBC (30 points each simulation). Reference equilibrium curve obtained using numerical continuation (- -). B) Time evolution of one simulation of $TetR$ and the control reference signal $TetR^*$ (–). C) Time evolution of $IPTG$ (i.e. control signal). Parameter values: $k_p = 0.0016$ and $aTc = 25ngmL^-1$.

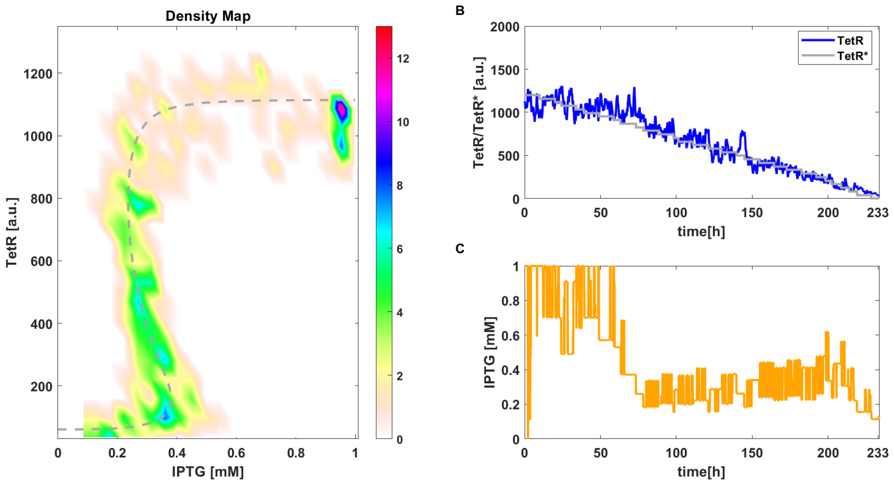


Figure S6: Short simulation (all points) of CBC with MPC applied to stochastic toggle switch model (4). A) Density plot of equilibrium curve measured using CBC (30 points each simulation). Reference equilibrium curve obtained using numerical continuation (- -). B) Time evolution of one simulation of $TetR$ and the control reference signal $TetR^*$ (–). C) Time evolution of $IPTG$ (i.e. control signal). Parameter values: $\gamma = 0.5$ (Eq. 25) and $aTc = 25ngmL^-1$.

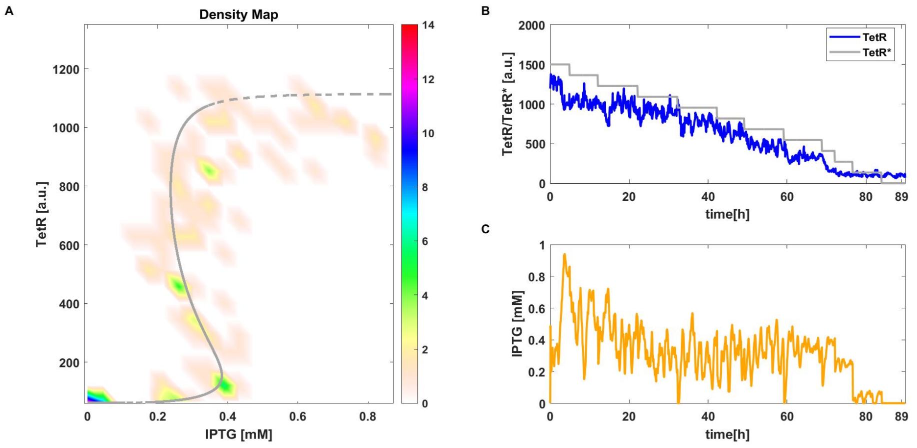


Figure S7: Short simulation of CBC with P controller applied to stochastic toggle switch model (4) in BSim. A) Density plot of equilibrium curve measured using CBC (12 points each simulation). Reference equilibrium curve obtained using numerical continuation (- -). B) Time evolution of one simulation of $TetR$ and the control reference signal $TetR^*$ (–). C) Time evolution of $IPTG$ (i.e. control signal). Parameter values: $k_p = 0.0016$ and $aTc = 25ngmL^-1$.

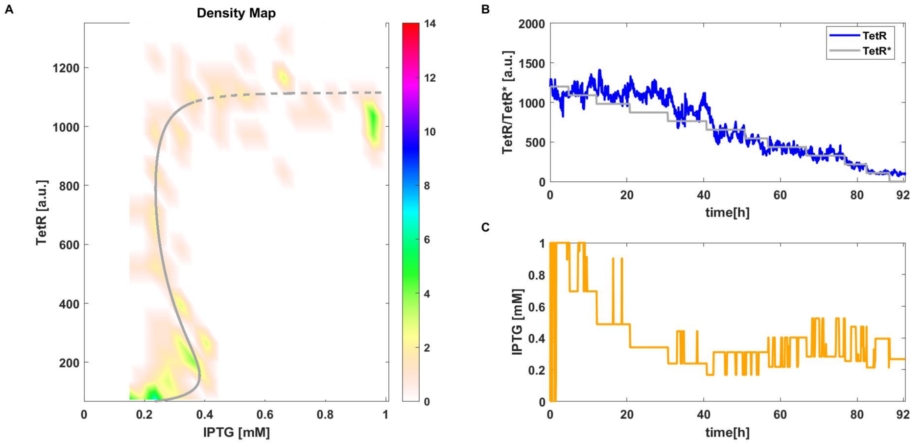


Figure S8: Short simulation of CBC with MPC applied to stochastic toggle switch model (4) in BSim. A) Density plot of equilibrium curve measured using CBC (12 points each simulation). Reference equilibrium curve obtained using numerical continuation (- -). B) Time evolution of one simulation of $TetR$ and the control reference signal $TetR^*$ (–). C) Time evolution of $IPTG$ (i.e. control signal). Parameter values: $\gamma = 0.3$ (Eq. 25) and $aTc = 25ngmL^-1$.

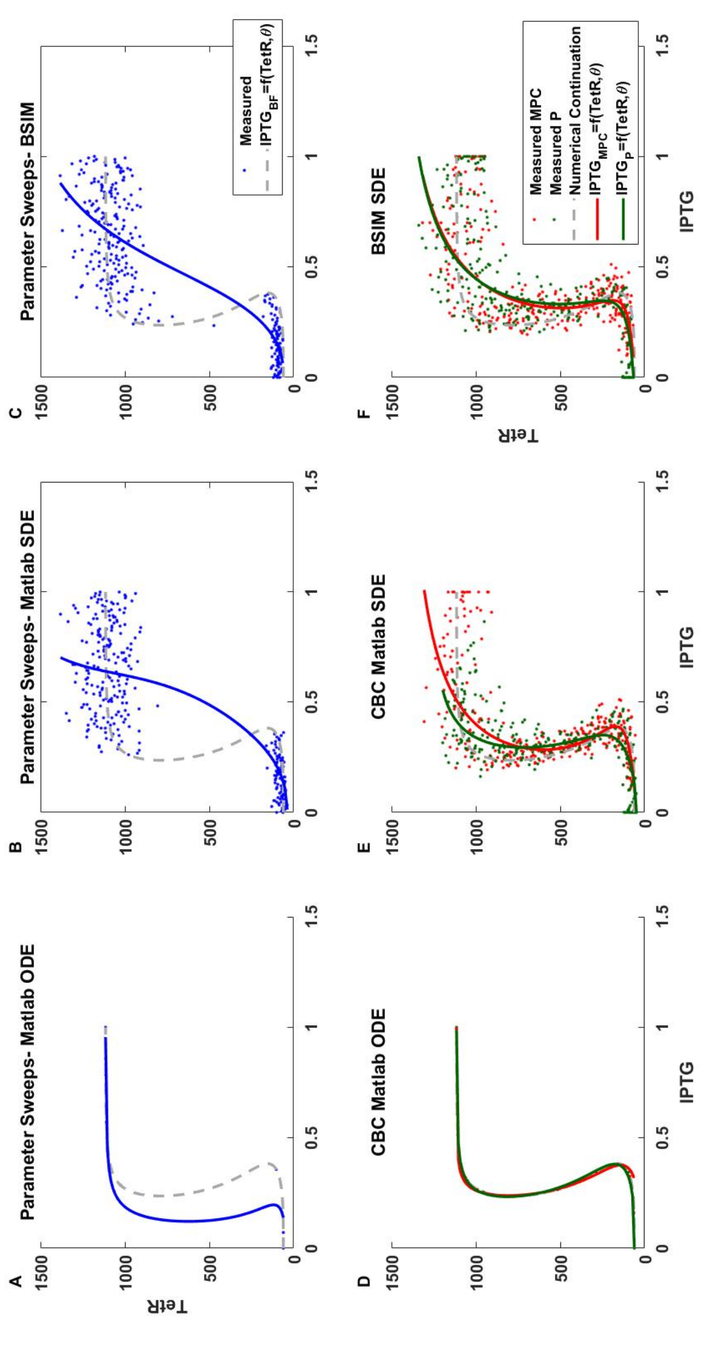


Figure S9: Caption next page

Figure S9: Equilibrium curve computed for model parameters estimated with constraints on IPTG complexity using parameter sweep data (–), CBC with MPC data (–) and CBC with P controller (–). Points collected with parameter sweeps (·), CBC with MPC (·) and CBC with proportional controller (·). A) Measured points from parameter sweeps in Matlab with the ODE toggle switch model and estimated curve. B) Measured points from parameter sweeps in Matlab with the SDE toggle switch model and estimated curve. C) Measured points from parameter sweeps in BSim with the SDE toggle switch model and estimated curve. D) Measured points from CBC routine in Matlab with the ODE toggle switch model and estimated curve. E) Measured points from the CBC routine in Matlab with the SDE toggle switch model and estimated curve. F) Measured points from BSim with the SDE toggle switch model and estimated curve.

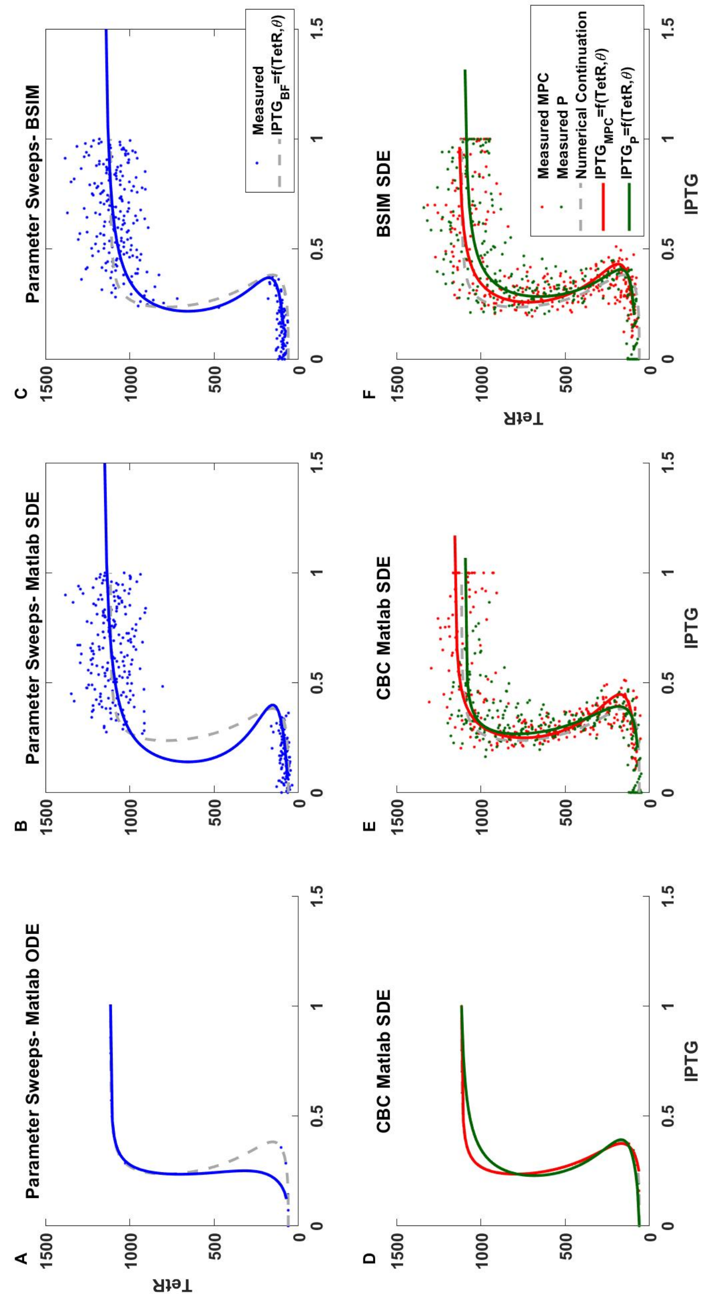


Figure S10: Caption next page

Figure S10: Equilibrium curve computed for model parameters estimated without constraints on IPTG complexity using parameter sweep data (—), CBC with MPC data (—) and CBC with P controller (—). Points collected with parameter sweeps (·), CBC with MPC (·) and CBC with proportional controller (·). A) Measured points from parameter sweeps in Matlab with the ODE toggle switch model and estimated curve. B) Measured points from parameter sweeps in Matlab with the SDE toggle switch model and estimated curve. C) Measured points from parameter sweeps in BSim with the SDE toggle switch model and estimated curve. D) Measured points from CBC routine in Matlab with the ODE toggle switch model and estimated curve. E) Measured points from the CBC routine in Matlab with the SDE toggle switch model and estimated curve. F) Measured points from BSim with the SDE toggle switch model and estimated curve.

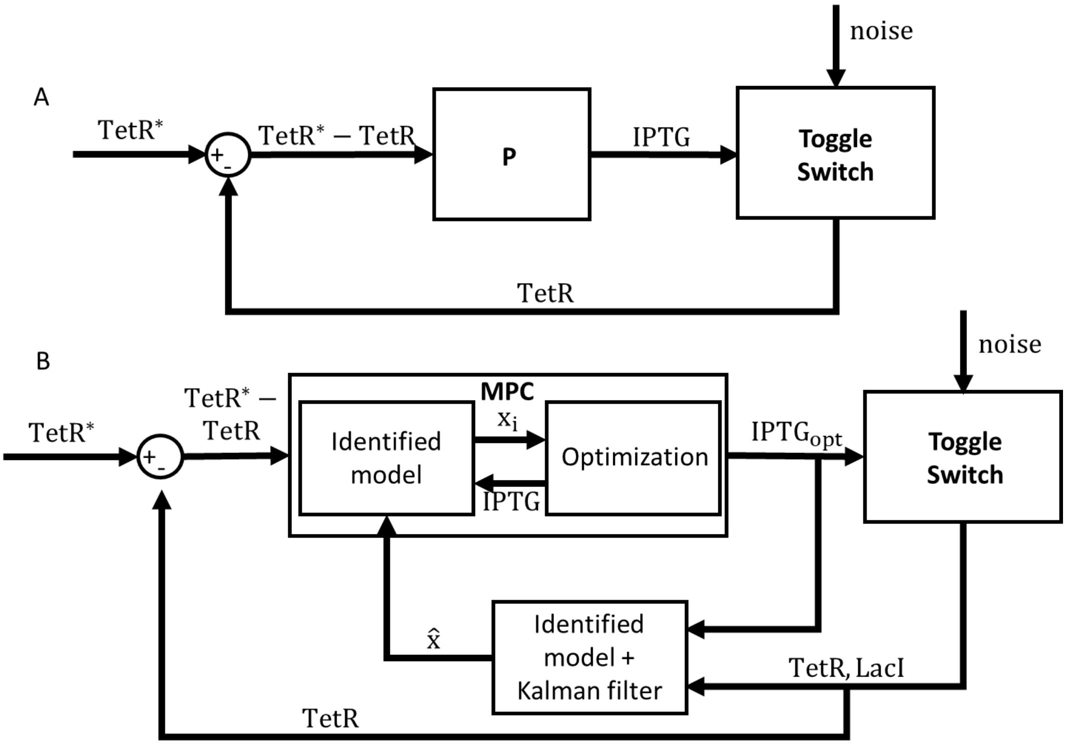


Figure S11: Control Feedback Loops implemented for CBC. A) Schematic of a proportional control loop. The measured system's output TetR is subtracted to the control reference signal $TetR^*$ to compute the error, that is fed to the controller. The output of the control block is the control signal IPTG, given to the toggle switch. B) Schematic of a MPC control loop and external Kalman filter for state estimation. $TetR^*$ is the control reference target, used to compute the error $TetR^* - TetR$. The MPC block is composed by the identified model, reproducing the state of the toggle switch $x_i$ over a certain prediction horizon, and the optimization block, that iteratively computes the control signal IPTG until the optimal $IPTG_{opt}$ is found. TetR and LacI are the toggle switch measured outputs and $\hat{x}$ is the estimated full state needed for the identified model.

| Parameter | Sweeps ODE | Sweeps SDE | MPC ODE | MPC SDE | P ODE | P SDE | BSIM MPC | BSIM P |
|---|---|---|---|---|---|---|---|---|
| $k_L^{m0}$ | 0.029 (-7.6%) | 0.016 (-50%) | 0.031 (-2.9%) | 0.048 (+50%) | 0.016 (-49.9%) | 0.048 (+50%) | 0.048 (+50%) | 0.048 +50%) |
| $k_T^{m0}$ | 0.076 (-35.3%) | 0.059 (-50%) | 0.125 (+5.8%) | 0.059 (-50%) | 0.093 (-21.4%) | 0.073 (-37.9%) | 0.094 (-20.4%) | 0.059 (-50%) |
| $k_L^m$ | 9.810 (18.2%) | 10.144 (22.2%) | 10.011 (20.6%) | 6.061 (-26.9%) | 4.468 (-46.1%) | 10.473 (26.1%) | 7.275 (-12.3%) | 10.9167 (-31.5%) |
| $k_T^m$ | 1.952 (-5.2%) | 3.089 (+50%) | 2.760 (+33.9%) | 2.819 (+36.8%) | 1.979 (-3.9%) | 2.575 (+25%) | 2.980 (+44.6%) | 2.952 (+43.3%) |
| $k_L^p$ | 0.960 (-1.2%) | 1.456 (+49.7%) | 1.059 (+8.9%) | 1.458 (+50%) | 0.911 (-6.2%) | 1.186 (+22%) | 1.458 (+50%) | 1.151 (+18.4%) |
| $k_T^p$ | 1.257 (+7.4%) | 1.261 (+7.8%) | 0.883 (-24.4%) | 1.461 (+24.9%) | 1.230 (+5.2%) | 1.284 (+9.8%) | 1.334 (+14%) | 1.170 (+0.05%) |
| $\theta_{LacI}$ | 23.356 (-26.8%) | 44.037 (+37.8%) | 24.988 (-21.7%) | 15.970 (-50%) | 47.255 (+47.9%) | 23.329 (-26.9%) | 15.970 (-50%) | 39.216 (+22.7%) |
| $\eta_{LacI}$ | 1.168 (-41.5%) | 1.157 (-42.1%) | 1.437 (-28.1%) | 1.000 (-49.9%) | 1.534 (-23.2%) | 1.000 (-49.9%) | 1.080 (-45.9%) | 1.000 (-49.9%) |
| $\theta_{IPTG}$ | 0.0812 (-10.2%) | 0.082 (-8.8%) | 0.096 (+6.7%) | 0.135 (+50%) | 0.135 (+50%) | 0.053 (-40.5%) | 0.135 (+50%) | 0.087 (-2.8%) |
| $\eta_{IPTG}$ | 2.655 (+32.8%) | 2.286 (+14.3%) | 2.500 (+25%) | 1.000 (-50%) | 2.133 (+6.6%) | 1.299 (-35%) | 1.000 (-50%) | 2.886 (+44.3%) |
| $\theta_{TetR}$ | 44.998 (+50%) | 43.130 (+43.7%) | 17.551 (-41.4%) | 18.918 (-36.9%) | 38.290 (+27.6%) | 20.830 (-30.5%) | 18.513 (-38.2%) | 44.999 (+50%) |
| $\eta_{TetR}$ | 2.919 (+45.9%) | 2.999 (+50%) | 2.806 (+40.3%) | 1.850 (-7.4%) | 2.999 (+49.9%) | 2.088 (+4.4%) | 1.714 (-14.2%) | 1.000 (-50%) |
| $\theta_{aTc}$ | 15.733 (+35%) | 6.906 (-40.7%) | 9.214 (-20.9%) | 16.547 (+42%) | 8.405 (-27.8%) | 12.218 (+4.8%) | 15.960 (+37%) | 5.825 (-50%) |
| $\eta_{aTc}$ | 2.999 (+50%) | 2.479 (+23.9%) | 2.446 (+22.3%) | 2.743 (+37.1%) | 1.771 (-11.4%) | 2.787 (+39.3%) | 1.636 (-18.1%) | 2.999 (+50%) |

Table S1: Caption next page.

Table S1: Estimated parameters for the toggle switch model depending on the method used to collect data. The first two columns show estimates computed after data collection via parameter sweeps. The other columns refer to different CBC experiments. For the deterministic simulation, estimates are computed out of 30 collected points, while for the stochastic simulations 300 points are considered. The percentage shown in brackets is the variation of the estimated parameter with respect to the original ones shown in table 1.

$$IPTG = \theta_{\mathrm{IPTG}} \left( \frac{ \frac{\mathrm{klm}\,\mathrm{klp}}{\left(-\frac{\mathrm{ktm}\,\mathrm{ktp}+\mathrm{ktm}_0\,\mathrm{ktp}-\mathrm{TetR}\,\mathrm{gtm}\,\mathrm{gtp}}{\mathrm{ktm}_0\,\mathrm{ktp}-\mathrm{TetR}\,\mathrm{gtm}\,\mathrm{gtp}}\right)^{1/\eta_{\mathrm{LacI}}}} - \mathrm{glm}\,\mathrm{glp}\,\theta_{\mathrm{LacI}} + \frac{\mathrm{klm}_0\,\mathrm{klp}}{\left(-\frac{\mathrm{ktm}\,\mathrm{ktp}+\mathrm{ktm}_0\,\mathrm{ktp}-\mathrm{TetR}\,\mathrm{gtm}\,\mathrm{gtp}}{\mathrm{ktm}_0\,\mathrm{ktp}-\mathrm{TetR}\,\mathrm{gtm}\,\mathrm{gtp}}\right)^{1/\eta_{\mathrm{LacI}}}} - \mathrm{glm}\,\mathrm{glp}\,\theta_{\mathrm{LacI}} \left(\frac{\mathrm{TetR}}{\theta_{\mathrm{TetR}}\left(\left(\frac{\mathrm{aTc}}{\theta_{\mathrm{aTc}}}\right)^{\eta_{\mathrm{aTc}}}+1\right)}\right)^{\eta_{\mathrm{TetR}}} + \frac{\mathrm{klm}_0\,\mathrm{klp}\left(\frac{\mathrm{TetR}}{\theta_{\mathrm{TetR}}\left(\left(\frac{\mathrm{aTc}}{\theta_{\mathrm{aTc}}}\right)^{\eta_{\mathrm{aTc}}}+1\right)}\right)^{\eta_{\mathrm{TetR}}}}{\left(-\frac{\mathrm{ktm}\,\mathrm{ktp}+\mathrm{ktm}_0\,\mathrm{ktp}-\mathrm{TetR}\,\mathrm{gtm}\,\mathrm{gtp}}{\mathrm{ktm}_0\,\mathrm{ktp}-\mathrm{TetR}\,\mathrm{gtm}\,\mathrm{gtp}}\right)^{1/\eta_{\mathrm{LacI}}}} }{ \mathrm{glm}\,\mathrm{glp}\,\theta_{\mathrm{LacI}}\left(\left(\frac{\mathrm{TetR}}{\theta_{\mathrm{TetR}}\left(\left(\frac{\mathrm{aTc}}{\theta_{\mathrm{aTc}}}\right)^{\eta_{\mathrm{aTc}}}+1\right)}\right)^{\eta_{\mathrm{TetR}}}+1\right) } \right)^{1/\eta_{\mathrm{IPTG}}} \tag{S1}$$

Movie S1: Simulation in time of CBC with P controller applied to deterministic toggle switch model (2) from Fig. 2. A) Equilibrium curve measured using CBC (·). $TetR(IPTG)$ transient trajectories (..). Reference equilibrium curve obtained using numerical continuation (- -). B) Time evolution of $TetR$ and the control reference signal $TetR^*$ (–). C) Time evolution of $IPTG$ (i.e. control signal).

Movie S2: Simulation in time of CBC with MP controller applied to deterministic toggle switch model (2) from Fig. 3. A) Equilibrium curve measured using CBC (·). $TetR(IPTG)$ transient trajectories (..). Reference equilibrium curve obtained using numerical continuation (- -). B) Time evolution of $TetR$ and the control reference signal $TetR^*$ (–). C) Time evolution of $IPTG$ (i.e. control signal).

Movie S3: Simulation in time of CBC with P controller applied to stochastic toggle switch model (4) in BSim. Only one simulation out of 10 from Fig. 6 is shown. Top left: microfluidic device with bacteria growth and division. Top right: time evolution of one simulation of $TetR$ (–) and the control reference signal $TetR^*$ (–). Bottom left: equilibrium curve measured using CBC (·). $TetR(IPTG)$ transient trajectories (–). Reference equilibrium curve obtained using numerical continuation (- -) Bottom right: time evolution of $IPTG$ (i.e. control signal).

Movie S4: Simulation in time of CBC with MP controller applied to deterministic toggle switch model (4) in BSim. Only one simulation out of 10 from Fig. 7 is shown. Top left: microfluidic device with bacteria growth and division. Top right: time evolution of one simulation of $TetR$ (–) and the control reference signal $TetR^*$ (–). Bottom left: equilibrium curve measured using CBC (·). $TetR(IPTG)$ transient trajectories (–). Reference equilibrium curve obtained using numerical continuation (- -) Bottom right: time evolution of $IPTG$ (i.e. control signal).

Movies S1, S2, S3 and S4 are attached as supplementary files.
